# Development of c-type lectin oriented surfaces for high avidity glycoconjugates: towards mimicking multivalent interactions on the cell surface

**DOI:** 10.1101/780452

**Authors:** Vanessa Porkolab, Carlo Pifferi, Ieva Sutkeviciute, Stefania Ordanini, Marwa Taouai, Michel Thepaut, Corinne Vivès, Mohammed Benazza, Anna Bernardi, Olivier Renaudet, Franck Fieschi

**Author notes:** These authors contributed equally (Vanessa Porkolab, Carlo Pifferi). CIC bioGUNE, Center for Cooperative Research in Biosciences, Bizkaia Science and Technology Park bld 801 A 48160 Derio, Bizkaia, Spain.

## Abstract

Multivalent interactions between complex carbohydrates and oligomeric C-type lectins govern a wide range of immune responses. Up to date, standard SPR (surface plasmon resonance) competitive assays have largely been to evaluate binding properties from monosaccharide units (low affinity, mM) to multivalent elemental antagonists (moderate affinity, µM). Herein, we report typical case-studies of SPR competitive assays showing that they underestimate the potency of glycoclusters to inhibit the interaction between DC-SIGN and immobilized glycoconjugates. This paper describes the design and implementation of a SPR direct interaction over DC-SIGN oriented surfaces, extendable to other C-type lectin surfaces as such Langerin. This setup provides a microscopic overview of intrinsic avidity generation emanating simultaneously from multivalent glycoclusters and from DC-SIGN tetramers that are organized in nanoclusters on the cell membrane. For this purpose, covalent biospecific capture of DC-SIGN via StreptagII /StrepTactin interaction offers the preservation of tetrameric DC-SIGN and the accessibility/functionality of all active sites. From the tested glycoclusters libraries, we demonstrated that the scaffold architecture, the valency and the glycomimetic-based ligand are crucial to reach nanomolar affinities for DC-SIGN. The glycocluster **3.D** illustrates the tightest binding partner in this set for a DC-SIGN surface (Kd= 18 nM). Moreover, the selectivity at monovalent scale of glycomimetic D can be easily analyzed at multivalent scale comparing its binding over different C-type lectin immobilized surfaces. This approach may give rise to novel insights into the multivalent binding mechanisms responsible to avidity and make a major contribution to the full characterization of the binding potency of promising specific and multivalent immunomodulators.

---

Pathogen recognition by the innate immunity system is governed by diverse interactions between immune system receptors, termed Pattern Recognition Receptors (PRRs), and pathogen-specific molecular structures called Pathogen-Associated Molecular Patterns (PAMPS). Although the vast majority of infections are prevented by the innate immunity, some of the pathogens have evolved to elude innate immunity components to evade or hijack the immune responses. A well-established example of such evasion mechanism is presented by DC-SIGN (Dendritic Cell (DC) specific ICAM-3 Grabbing Non-integrin), a PRR abundantly expressed on immature DCs, which patrol peripheral tissues for invading pathogens and are key players in the activation of adaptive immune responses. (1, 2) Multiple studies have indicated that a wide range of pathogens (such as HIV, Dengue, HCMV, M. tuberculosis and others) uses DC-SIGN as a port for infection, promotion and dissemination. (2–4) These findings have placed DC-SIGN as an important therapeutic target and many efforts are being invested to develop DC-SIGN antagonists. (5–13)

DC-SIGN is a tetrameric C-type lectin receptor (CLR) with specificity to D-mannoseand L-fucosecontaining oligosaccharides that bind the C-terminal carbohydrate recognition domain (CRD) in a Ca2+-dependent manner. (1, 14) Although the intrinsic affinity of a single CRD for a monosaccharide is low (KD in mM range), the global DC-SIGN/multivalent carbohydrates interaction affinity is markedly amplified (KD from µM to pM) through the avidity phenomenon.(15, 16) The latter is obtained thanks to the combined multivalent presentation of the glycan from the pathogen on one side and the CRD due to DC-SIGN tetramerization on the other. In addition, a specific enhancement of the avidity is correlated to the concentration of DC-SIGN into lipid rafts on the cell membrane. (17, 18) Such density of DC-SIGN binding sites within small two-dimensional areas increase the interaction strength by multiplying binding events.

The multivalent nature of DC-SIGN interaction with pathogens has dictated the general strategy for the design of DC-SIGN antagonists. First, efforts have been made towards the development of glycomimetic compounds, with an improved affinity and specificity, interacting with single CRD moities. Secondly, different polyvalent scaffolds displaying glycomimetics should be also considered with care, considering the various modes of avidity that can occur at the cell surface (Figure 1A). (19, 20)

**Figure 1:**
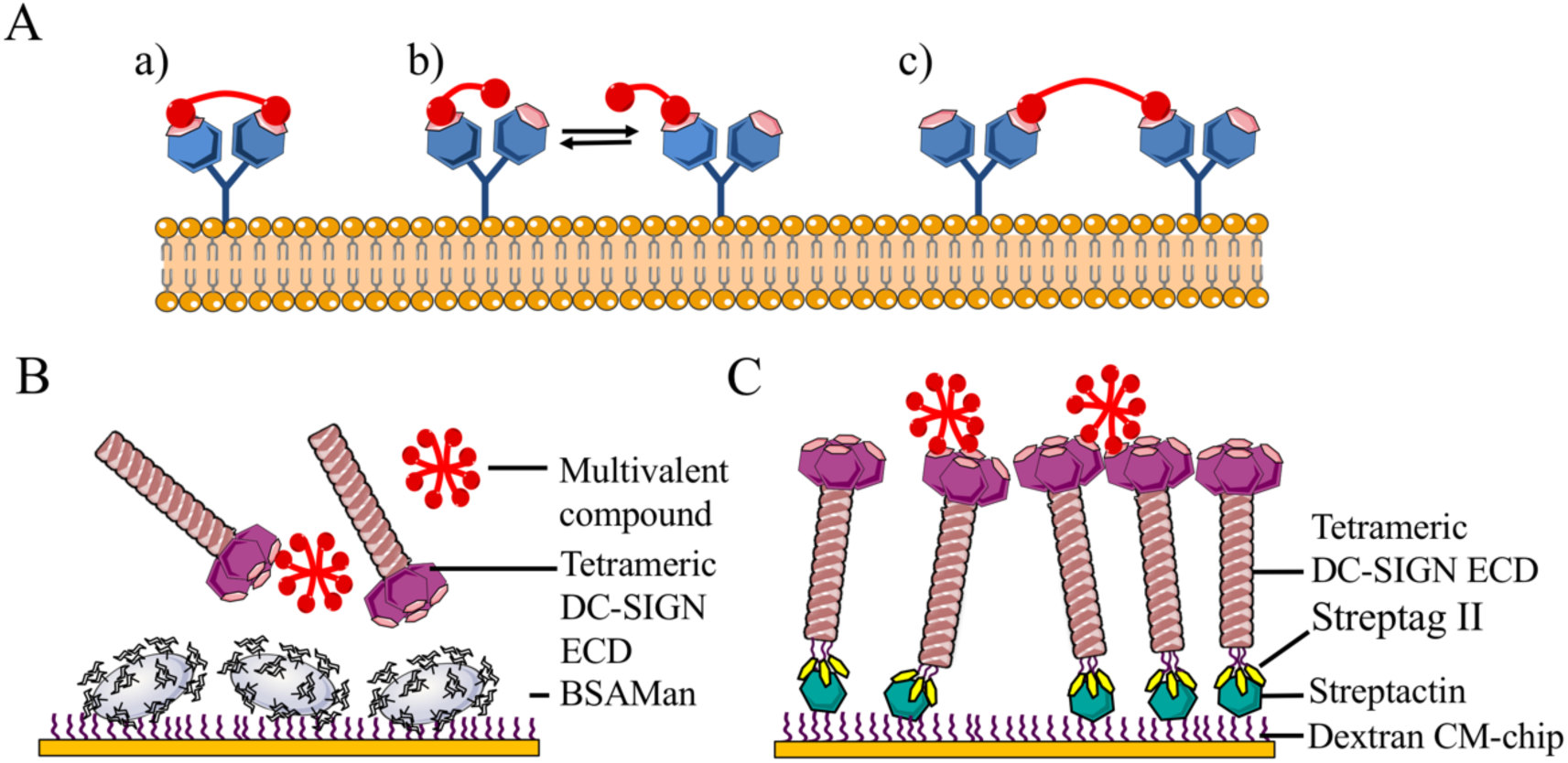
Multivalent binding of DC-SIGN. (A) Identification of potential binding modes between a bivalent ligand (red spheres) and two binding sites at cell surface. The chelating effect (a), the statistical rebinding (b), the clustering effect contributes to the avidity improvement. (B) Principle of DC-SIGN competition assay. Man-BSA is covalently immobilized onto the CM surface. Increasing concentrations of multivalent compound are co-injected with a given concentration of DC-SIGN ECD. (C) DC-SIGN S-ECD is immobilized by its N-terminal *Strep*tagII tag onto a *Strep*Tactin-functionalized CM3 sensor chip. Increasing concentrations of multivalent compound are injected over the surface.

Indeed, the rational development of compounds with high avidity requires a method with experimental set ups that enables all compound binding modes and allows affinity determination in nM-pM range. Surface plasmon resonance (SPR) allows label-free real-time interaction studies for flexible assay design. Many groups, including ours, use an SPR-based competition assay for evaluation of both monovalent and polyvalent compounds as DC-SIGN binders (Figure 1). Although screening by indirect competition assay in most cases is sufficient to identify monovalent lead compounds and first generation of multivalent structures, optimized multivalent compounds require direct interaction studies. Indeed, while the competition assay indirectly allows evaluation of low-moderate affinity, it does not provide information on direct properties of the complex, such as the association rate or the complex stability, in which the tested compound is involved (Figure 1B). Moreover, IC50 value cannot be measured for compounds with an affinity for DC-SIGN higher than that of the reporter system typically used in the competition assays (DC-SIGN/BSA-Man surface). (8) To overcome these limits of the competition assay, monitoring the direct interaction between multivalent compounds and a DC-SIGN surface would be the best approach to evaluate surface-generated avidity (Figure 1C). (8, 21)

To develop a direct interaction method that in some way mimics the cell surface, DC-SIGN has to be immobilized covalently and unidirectionally over the surface. Although direct interaction of various ligands with DC-SIGN surface has been already examined, (22–24) in all these studies the tetrameric DC-SIGN ECD was immobilized via the standard amino coupling procedure, which requires preparation of the protein in acidic condition. Even though the prepared surfaces retain sugar-binding activity, CRDs are sensitive to acidic conditions (25, 26) and moreover the tetrameric organization of DC-SIGN is disrupted due to dissociation into monomers induced by acidification. (26) Consequently,, the binding activity observed does not necessarily reproduce the native interactions features with CRDs presentation as tetramers with a correct topology. In addition, the randomly immobilized DC-SIGN does not exactly reflect the biological accessibility of the lectin, which is anchored to the cell membrane through its neck region. An uncontrolled non oriented immobilization of DC-SIGN on the surface, may decrease the CRD accessibility for multivalent ligands, thus disturbing the interaction process.

In this paper, we first describe a rigorous approach of DC-SIGN surface preparation that could be extended to other C-type lectins, such as Langerin. The bio-specific capture of DC-SIGN, in a buffer at physiological pH yields undisturbed tetrameric DC-SIGN ECD surfaces with well-oriented CRDs. In that setting, we are able to accurately estimate the affinity and the avidity of multivalent optimized compounds. To exemplify the relevance of this method, the binding properties of selected optimized glycoconjugates (glycoproteins and different series of glycoclusters) were investigated and the results compared with the standard SPR inhibition assay. In such way, this direct interaction assay with oriented C-type lectin immobilization constitutes a further step toward the characterization of the multivalent binding complexity.

## RESULTS AND DISCUSSION

### Development and characterization of DC-SIGN S-ECD surface

In order to prepare an intact tetrameric DC-SIGN ECD oriented surface, we have chosen to use the biospecific capturing approach. DC-SIGN ECD construct tagged at its N-terminus with a *Strep*tagII (here after called DC-SIGN S-ECD) is captured by the immobilized *Strep*Tactin over a CM dextran sensor chip (Figure 1C). For the initial estimation of capturing stability, the affinity of DC-SIGN S-ECD to *Strep*Tactin was determined by the titration of covalently immobilized *Strep*Tactin surface with DC-SIGN SECD (Figure S22). Unfortunately, despite a relatively high affinity (57 nM), DC-SIGN S-ECD surface is not stable enough and DCSIGN dissociation from *Strep*Tactin surface was observed from cycle to cycle (data not shown).

In order to overcome this problem, covalent stabilization of captured DC-SIGN S-ECD was probed. To this end, after covalently immobilizing *Strep*-Tactin^®^ on the dextran CM3 surface, DC-SIGN S-ECD in pH 7.4 buffer was injected after a re-activation of the dextran/*Strep*Tactin surface by EDC/NHS mixture (Figure 2A). To control the binding specificity, DC-SIGN ECD, *i.e.* a construct without *Strep*tagII was also injected over reactivated dextran/*Strep*Tactin surface. No binding was detected for the latter (Figure 2B). Indeed, DC-SIGN ECD at pH 7.4 is largely negatively charged (pI = 5.16) and is repulsed from the negatively charge surface of the sensor preventing its functionalization. It confirms that it is the biospecific interaction between the StreptagII, at the N-ter of the S-ECD construct, and *Strep*Tactin that preconcentrate the protein at the surface allowing its reaction with activated groups. DC-SIGN S-ECD is indeed immobilized through the N-terminus of DC-SIGN ECD in a well-oriented manner. Moreover, the oligomeric structure integrity was preserved through the immobilization of DC-SIGN S-ECD at physiological pH.

**Figure 2.**
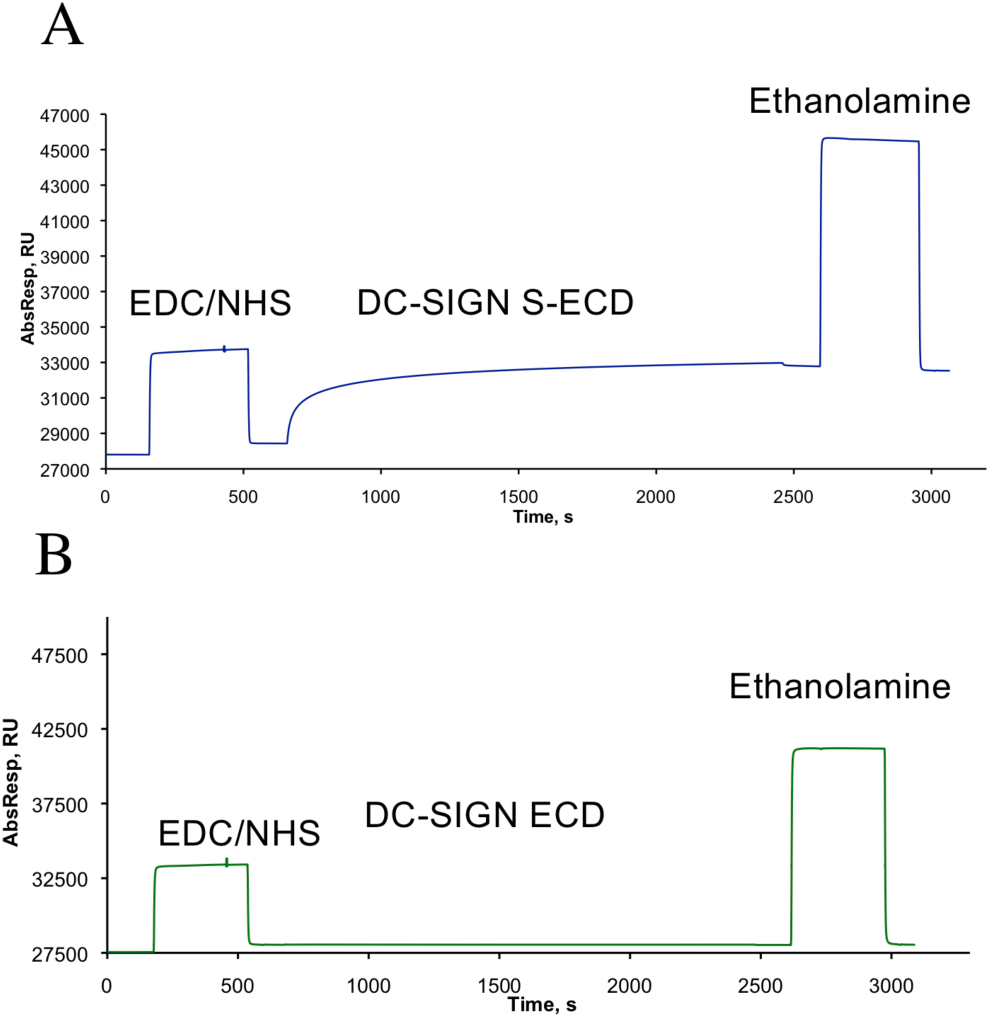
The sensorgrams showing the immobilization of DC-SIGN constructs on reactivated dextran/*Strep*Tactin surface. The dextran/*Strep*Tactin surface was reactivated by injection of EDC/NHS mixture. Then DC-SIGN S-ECD (A) or DC-SIGN ECD (B) in HBS-P running buffer were injected over reactivated surface. The remaining activated -COOH groups were blocked by ethanolamine. More details about the protocol are available in SI.

Covalent stabilization of the soft DC-SIGN S-ECD bio-capture resulted in optimal activity and stability of the surface, as illustrated on Figure 3A. Repetitive injections of BSA-Man glycoprotein enable to clearly to attest the maintenance of the surface activity and stability. So far, the immobilization strategy was focused on the well-oriented, stable and functional DC-SIGN ECD over a CM surface, the following section presents the binding characterization of multivalent glycoconjugates over the DC-SIGN S-ECD surface.

**Figure 3.**
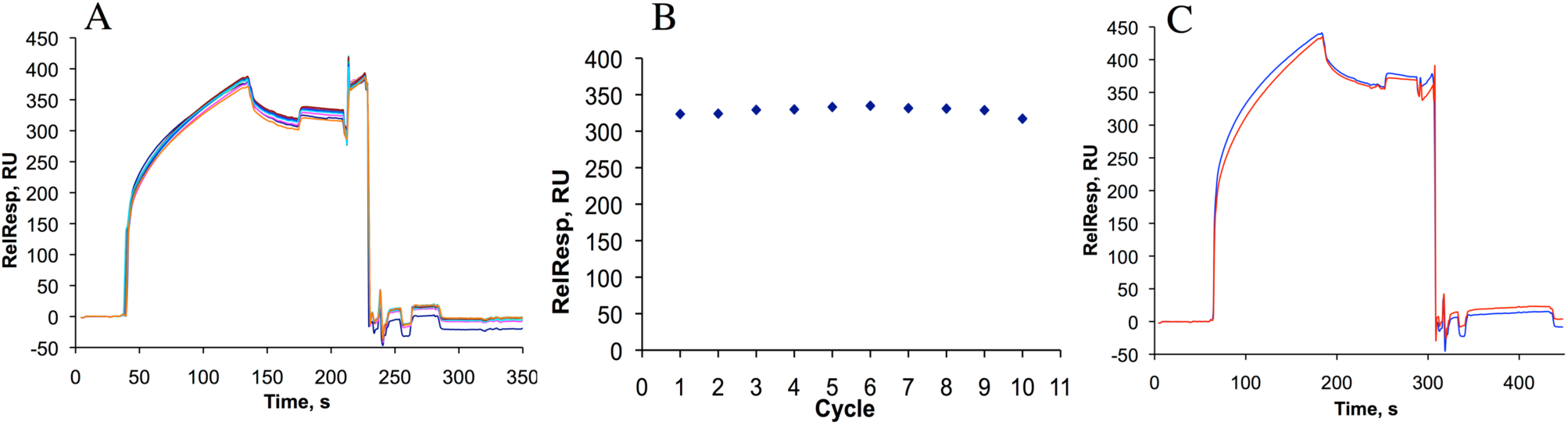
The evaluation of DC-SIGN surface stability. (A) Reference surface corrected overlaid sensorgrams showing 10 consecutive injections of BSA-Man over DC-SIGN S-ECD surface. Each injection was followed by a regeneration step. (B) The corresponding binding responses measured after each injection and plotted against cycle number (*i.e.* injection number). (C) Reference surface-corrected sensorgrams showing the first (blue) and the last (red) BSA-Man injection controls of a series of 44 cycles.

### Comparison of BSA-Man / DC-SIGN ECD interaction analyzed in the competitive or direct interaction set up

Before studying the interactions between the DC-SIGN-oriented surface and a small library of glycoconjugates with different valencies and spatial architectures, the interaction with glycoprotein BSA-Man bearing 12 glycosylated sites has been first characterized by direct interaction assay. Figure 4 compares DC-SIGN titration toward a BSA-Man surface (Figure 4A, corresponding to the competition assay set up) and the BSA-Man titration over a DC-SIGN S-ECD oriented surface (Figure 4B). The sensorgram shape and the kinetics of association look completely different between the two settings (compare Figure 4A(a) and 4B(a)). In the latest, the association phases apparently consist of unique fast portion for the lowest concentrations and an early fast portion followed by a slow portion for higher concentrations.

**Figure 4.**
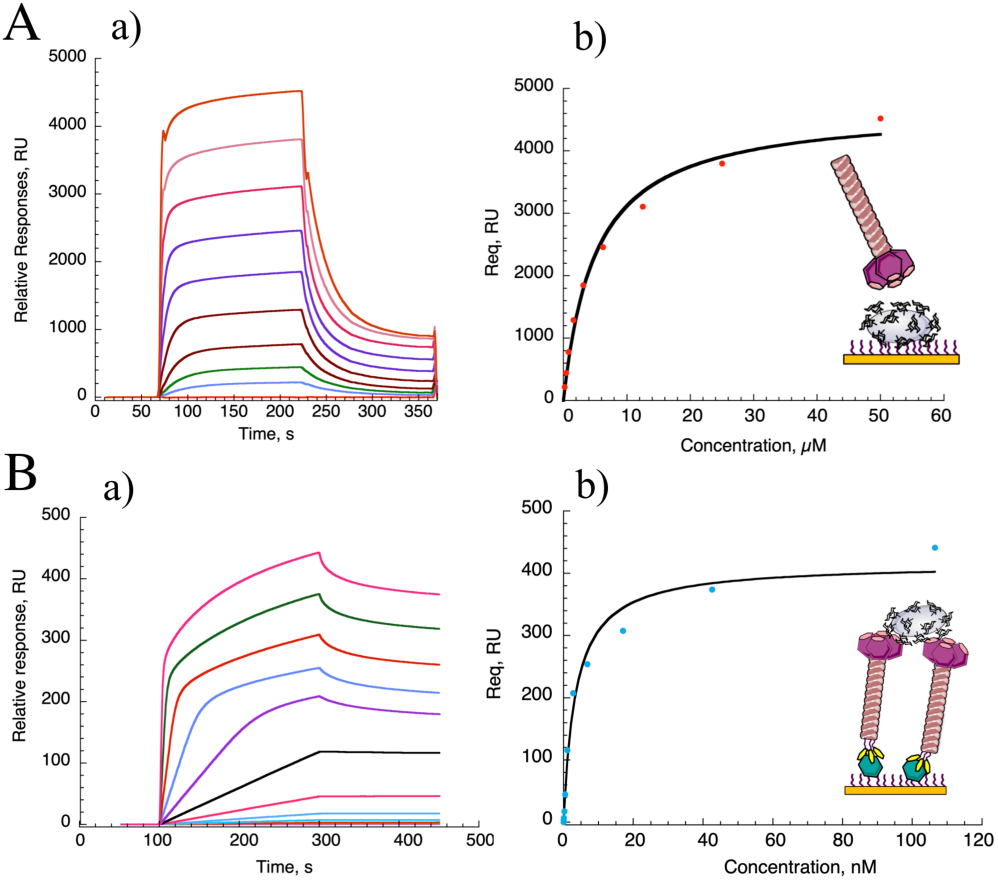
Interaction of BSA-Man with DC-SIGN ECD. (A) DC-SIGN ECD titration over a BSA-Man surface (1600 RU) (a) Reference surface corrected sensorgram (b) titration curve obtained by plotting binding responses against DC-SIGN ECD concentration (red plot). (B) BSA-Man titration over a DC-SIGN S-ECD oriented surface (2500 RU). (a) Reference surface corrected sensorgram and (b) titration curve obtained by plotting binding responses against BSA-Man concentration (blue plot). The two fits (black curve in (b)) were obtained by a *steady state affinity* model.

The coexistence of different multivalent binding modes (Figure 1A) could be translated as different kinetics and may cause this phenomenon. Moreover, the dissociation phase was markedly slower on DC-SIGN S-ECD surface suggesting an improvement of multivalency. In the first case (Figure 4A), where DC-SIGN ECD is the analyte, a maximum of 4 simultaneous binding events by tetramer can theoretically occur with the surface, while in the second set-up, BSA-Man displaying up to 12 glycans can establish up to 12 theoretical binding events with the oriented surface (probably less, but still higher than 4). Regarding the affinity estimation, the complexity of the binding event led us to investigate different analytical methods to have access to a multivalent KD value associated to the surface (K_D apparent_). These analyses are described in supplementary materials. Here, the most relevant and equitable method is the *steady state affinity* model. Even if the stoichiometric ratio is not necessarily respected during the association phase, we assumed than the overall and average binding is close to simple Langmuirian 1:1 binding model considering the surface as a whole as being the ligand.

Thus, DC-SIGN ECD titration over a BSA-Man surface (Figure 4A) predicts a K_D app_ equal to 3.74 µM, while BSA-Man titration onto an oriented DC-SIGN S-ECD surface (Figure 4B) gave an affinity associated to the avidity surface (K_D apparent_) of 5 nM. In spite of the fact that characterization of multivalent binding mechanisms are not accessible, the development of oriented DC-SIGN S-ECD surface highlights the improvement of avidity generated by the immobilization of receptors onto a surface. It maximizes the binding of glycoconjugates as it is expected to occur at the cell surface. Statistically, the combination of multivalent binding events is increased when the partner with the lowest number of binding sites (*i.e.* 4 CRDs of DC-SIGN) is coated onto the surface while the analyte with the highest number of binding unit (*i.e.* 12 sites of glycosylation of BSA) is free in solution. This suggests that the avidity of multivalent compounds might be underestimated if they are only analyzed in a standard competition assay. From these findings, the oriented surface proved its relevance and should become the standard method to measure the affinity of rationally designed multivalent glycoconjugates.

### SPR binding analysis of DC-SIGN ECD/ glycoclusters

The design of glycoconjugates targeting DC-SIGN aims to produce an avidity comparable to or better than what happens at the cell surface. Here, promising multivalent compounds have been designed dealing with different geometries and spatial organizations. These efforts have been made in order to consolidate the binding with DC-SIGN by promoting chelating, rebinding and clustering binding events. We describe herein the interaction study of DC-SIGN with glycoclusters bearing multiple copies of mannose residues (ligand **A**) or optimized glycomimetics, previously described by our laboratory (ligands **B, C, D**) (12, 27–29) (Figure 5).

**Figure 5.**
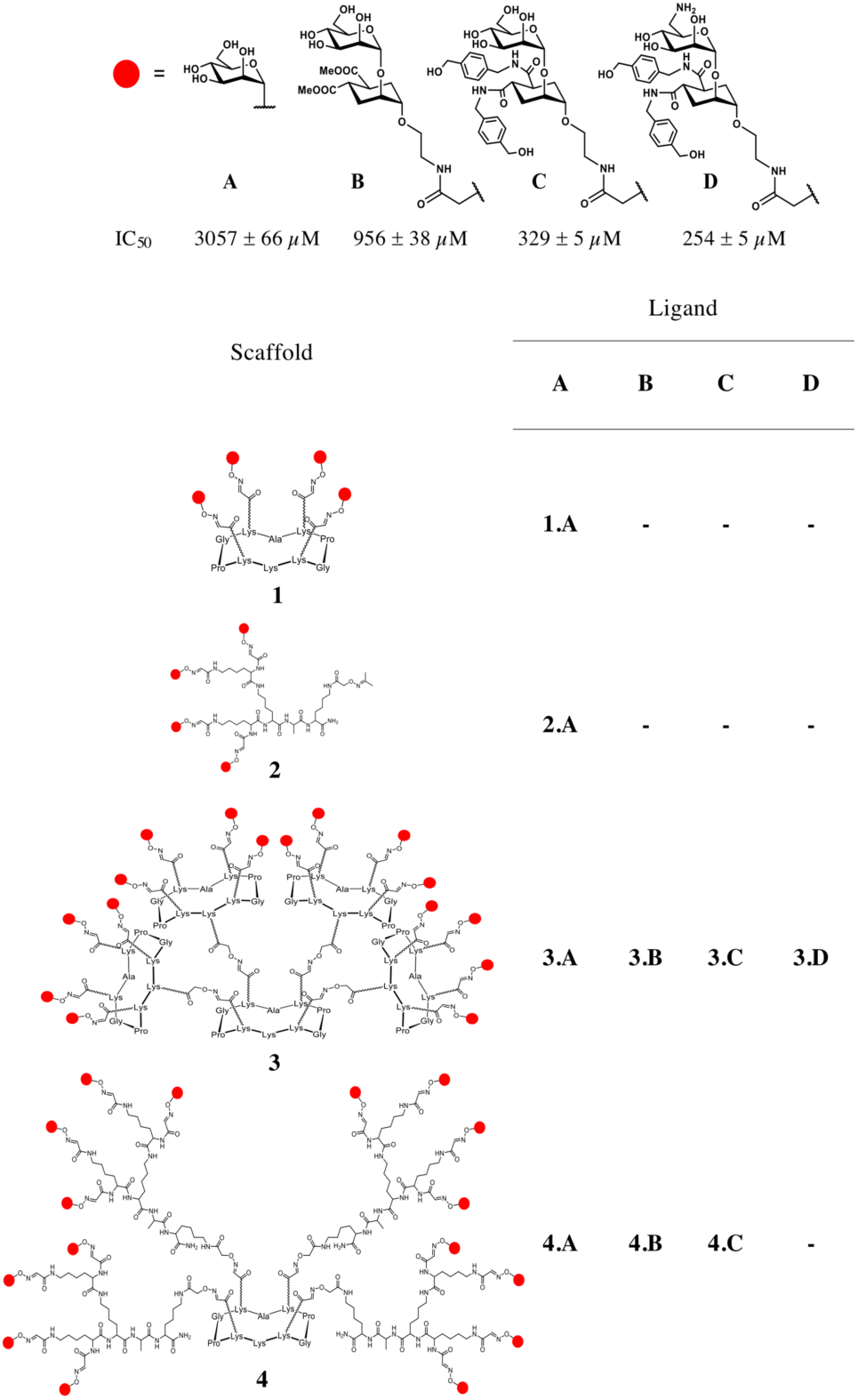
Structures of tested glycoclusters. Ligands (red spheres) are represented at the upper panel with their corresponding relative affinity (IC_50_) for DC-SIGN previously reported in SPR competition experiments. (12)

Supramolecular scaffolds **3** and **4** both display a valency of **16** ligands, but differs in the manner the ligand is displayed. Although bearing the same core platform, scaffold **3** is equipped with four rigid tetravalent display units, while peripheral arms of scaffold **4** allow for a higher degree of flexibility. The panel of glycoclusters described herein was generated by conjugating aldehyde-bearing scaffolds to aminooxy-functionalized carbohydrate ligands through oxime ligation (see Supporting Information). The relative affinity of these compounds towards DC-SIGN were first analyzed, with the competitive assay set-up, for their ability to inhibit the binding of DC-SIGN ECD to BSA-Man immobilized surface (Figure 1B). Increasing concentrations of scaffold **3** and **4**-based compounds were co-injected with 20 µM of DC-SIGN ECD. The sensorgrams and inhibition curves are shown on the supplementary materials (Figure S24 and S25) and relative affinity results (IC_50_ values) are compared in Figure 6A. As shown in Table 1, all compounds have an IC_50_ comprised between 2 and 5 µM. The first observation indicates that the affinity does not critically varies with the scaffold choice nor the ligand optimization (from A to C). However, the IC_50_ values are situated in the same µM affinity range than the reporter system (DC-SIGN ECD/ BSA-Man-surface, see Figure 4A), thus the results suggest that the intrinsical limit of the inhibition assay has been reached.

**Table 1:** IC_50_ and *K*D values for glycoclusters.

| Compounds | Valency | $\text{IC}_{50}$, $\mu\text{M}$ | $K_{\text{D app}}$, $\mu\text{M}$ |
| --- | --- | --- | --- |
| **1.A** | 4 | $1009 \pm 10.4$ | - |
| **2.A** | 4 | $129 \pm 4.7$ | - |
| **3.A** | 16 | $3.0 \pm 0.03$ | $6.3 \pm 1.90$ |
| **3.B** | 16 | $2.8 \pm 0.04$ | $1.37 \pm 0.15$ |
| **3.C** | 16 | $4.9 \pm 0.05$ | $1.11 \pm 0.12$ |
| **4.A** | 16 | $2.5 \pm 0.02$ | $2.11 \pm 0.22$ |
| **4.B** | 16 | $2.3 \pm 0.01$ | $0.78 \pm 0.03$ |
| **4.C** | 16 | $4.4 \pm 0.06$ | $0.38 \pm 0.02$ |

**Figure 6.**
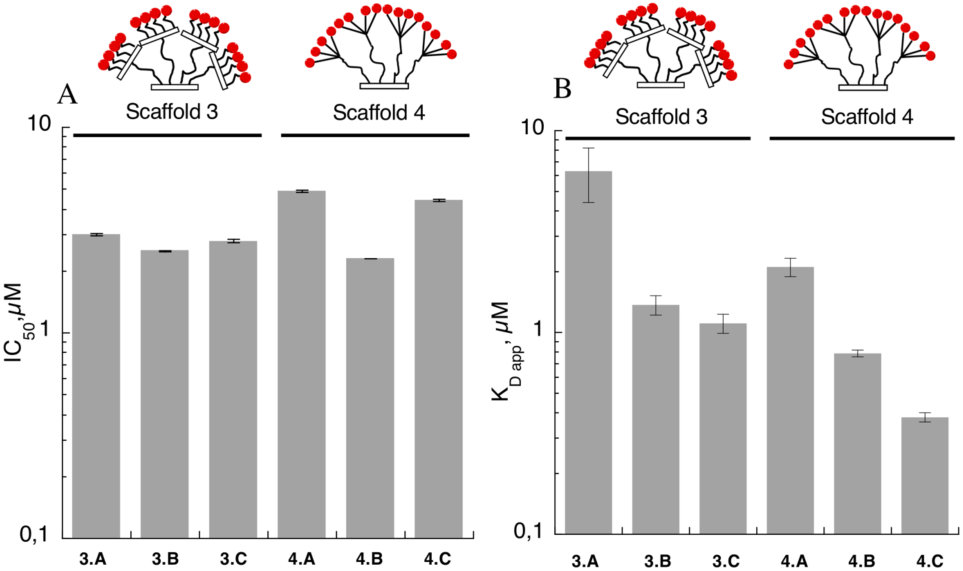
Affinities of glycoclusters for DC-SIGN (A) Comparison of IC50 values obtained by SPR competition assays. (B) Comparison of K_D app_ values over a DC-SIGN S-ECD oriented surface.

These results further underline the limitation of the reporter system limitation. For glycoconjugates with affinities higher than that of the reference IC_50_ values cannot be determined. This study clearly shows that SPR competition assays with BSA-Man is an useful tool to evaluate relative affinities for low-moderate affinity compounds. Nevertheless, for compounds with tight affinities, the indirect method is misleading, since in the multivalent binding mode context, in which the absolute affinity is mostly driven by the avidity.

To increase avidity, compounds are then tested by direct interaction. Increasing concentrations of glycoconjugates were injected over a DC-SIGN S-ECD oriented surface and then binding responses were plotted against compound concentration (Figure S26). The sensorgrams have the same trend as the BSA-Man titration over a DC-SIGN S-ECD oriented surface (Figure 4B). Thereby, the different kinetic behaviors during the association phase and the slow dissociation phase confirm that the binding is governed by multivalent interactions. The observed K_D app_ values of scaffold **3** and **4** displaying glycomimetics **A, B** and **C** are collected in Table 1 and show that the combination of scaffold **4** with ligand C corresponds to the best affinity for a DC-SIGN S-ECD surface (K_D app_ =0.38 ± 0.02 µM).

Moreover, the optimization of terminal ligand from **A** to **C** plays an important role in affinity improvement and could increase up to 6-fold the binding (from K_D app_= 6.3 µM for **3.A** to 1.11 µM for **3.C** and from 2.11 µM for **4.A** to 0.38 µM for **4.C**).

The comparison of apparent affinities (Figure 6B) shows the following tendency: the flexible extremities of dendrimers (scaffold **4**) enhance the affinity for DC-SIGN surface compared to terminal rigid display units. Comparing both scaffold displaying 16 copies of ligand C, the dendrimer-scaffold in **4.C** (K_D app_=0.38 µM) confers a 3-times improvement of affinity than the scaffold in **3.C** (K_D app_=1.1 µM). Therefore, the affinity of the optimized ligand and the ability of the scaffold to generate multivalency are two parameters impacting the avidity toward a DC-SIGN S-ECD surface. The flexibility of dendrimers **4** seems to enhance the combination of multivalent binding modes and, *inter alia*, can simultaneously bridge simultaneously two DC-SIGN active sites resulting in a nM range affinity. Indeed, maximal distances between sugar moieties have been previously estimated to be between 80 −70 Å for scaffold 3 and 4. (27)

Overall, these results suggest that the oriented DC-SIGN S-ECD surface allows to evaluate the K_D app_ of multivalent high affinity compounds, not measurable by a standard SPR competition assay. Further deeper investigations can be now carried out to evaluate the contribution of the ligand and the scaffold in the overall binding of a multivalent compound over a DC-SIGN S-ECD surface.

### Binding and selectivity investigation of the glycocluster 3.D over a DC-SIGN S-ECD surface

Working with different scaffolds displaying several ligands could interfere or increase the interaction with the biological target. Through this experiment, the nonspecific interaction from the scaffold will be estimated. For that observation, compound **3.0**, the common precursor (27) of glycoclusters **3.A-D**, devoid of the carbohydrate moiety, was selected for its high solubility in buffer solution (see Scheme S2). Furthermore, scaffold **3** was also functionalized with 16 copies of glycomimetics **D** that bears with a positive charge due to amine protonation. This monovalent ligand is known to bind DC-SIGN selectively vs Langerin, another C-type lectin receptor of similar specificities. (12) The selectivity of cluster **3.D** toward DC-SIGN will be also analyzed, in the further step, comparing the two lectin immobilized surfaces *i.e.* DC-SIGN S-ECD and Langerin S-ECD surfaces.

The titration of cluster **3.D** over the DC-SIGN oriented surface shows the different kinetics for each injection that clearly illustrate the cumulative multivalent binding that occur between cluster **3.D** and DC-SIGN S-ECD surface. From the steady-state binding analysis (Figure 7A), the cluster **3.D** appears to be a highly efficient ligand for DC-SIGN S-ECD surface with a K_D app_ equal to 18 nM. Moreover, the substitution of ligand **C** by **D** dramatically increases the affinity for DC-SIGN (1 µM for **3.C**, 18 nM for **3.D**).

**Figure 7:**
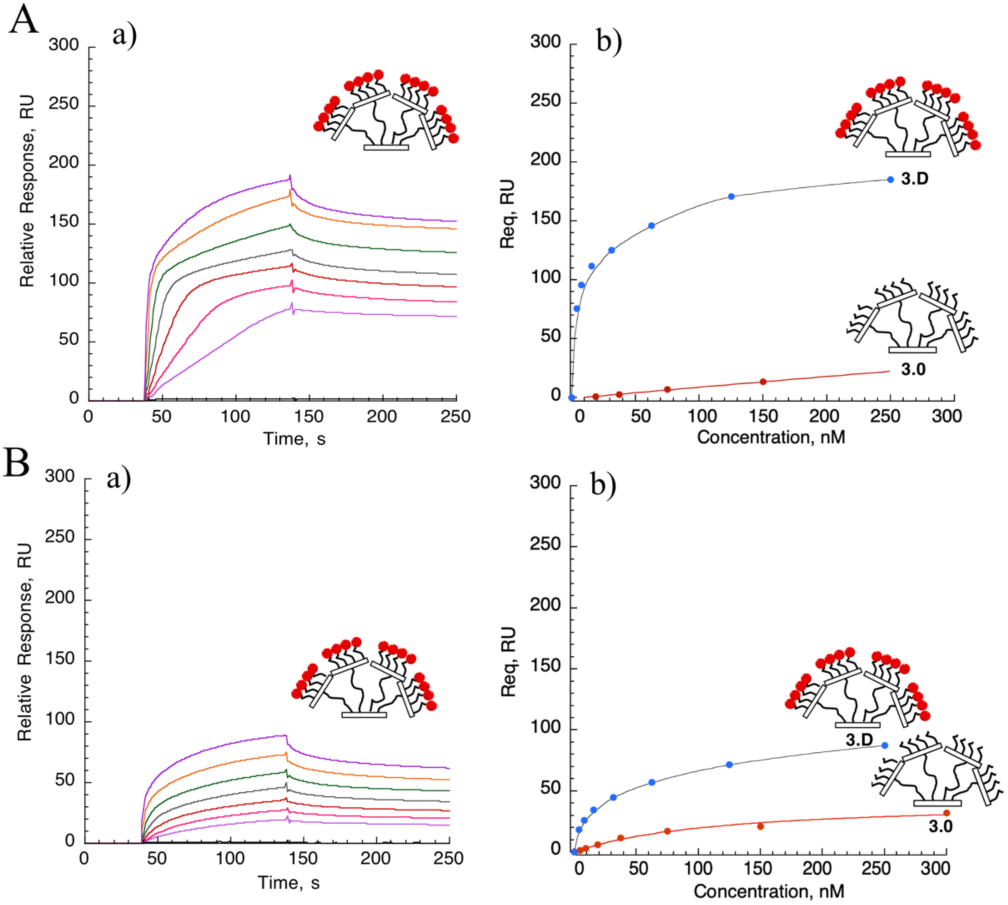
Titration of cluster **3.D** and its scaffold **3.0** over DC-SIGN S-ECD oriented surface (A) and Langerin S-ECD oriented surface (B). (a) Reference-subtracted sensorgram of increasing concentrations (0, 4, 8, 16, 31, 62, 125, 2_50_ nM) of 3.D. (b) Steady state binding analysis (n=1) for **3.D** (blue dots). Linear regression and steady state fit are respectively applied for scaffold **3.0** (red dots) over a DC-SIGN S-ECD oriented surface (A) and Langerin S-ECD oriented surface (B).

In order to know if the binding potency of **3.D** is fully active-site directed, the naked scaffold **3.0** was also titrated over the DC-SIGN S-ECD surface. The Req signals, subtracted by reference surface and running buffer, were plotted against the scaffold **3.0** concentrations and fitted by linear regression (Figure 7A (b)). The data highlight the slight non-specific binding of scaffold **3.0** over the surface. For instance, comparing both scaffold **3.0** and **3.D** at 250 nM, the nonspecific binding contribution corresponds to 8 % of the overall binding. According to these data, we can infer that i) the contribution of the naked scaffold **3.0** is considered as irrelevant regarding to the overall binding and ii) the multivalent displaying of ligand **D** (18 nM) significantly improves the affinity for DC-SIGN compared to its counterpart **3.C** (1 µM).

Another significant aspect of ligand **D**, compared to ligand **B** and **C**, is its absolute selectivity toward DC-SIGN. It has been demonstrated by SPR competition assays that the glycomimetic **D** interacts with DC-SIGN ECD and no interaction were measurable with Langerin ECD, impairing any selectivity factor evaluation. (12) However, up to now, the selectivity studies have only been carried out at the monovalent scale. As expected, an increase of affinity from monovalent to multivalent displaying of glycomimetics **D** can be reached through a massive avidity effect generated by the multivalent glycocluster **3.D** and the oriented DCSIGN S-ECD surface. It may be possible that the polycationic nature of **3.D** brings supplementary electrostatic, with the electronegative DC-SIGN active site, that amplify its avidity effect. Avidity is a well-known fundamental property able to improve the affinity of multivalent interactions over the monovalent scale, however, it is not yet clear whether avidity would preserve selectivity. Here, we explore the selectivity effect of cluster **3.D** over a Langerin S-ECD surface. The results obtained from the steady state binding analysis of cluster **3.D** and its corresponding scaffold **3.0** are presented on Figure 7B.

The cluster **3.D** strongly interacts with the Langerin S-ECD surface and the KD app extracted from the fit is equal to 205 nM. However, during the association phase, no significant difference of kinetic is observed. A possible explanation might be that the interaction between the compound and the surface does not only result from a a multivalent binding association but that another type of interaction is also engaged. These observations raise the possibility of non-specific interactions from the empty scaffold **3.0** or the glycomimetic **D** itself. The titration of **3.0** over the Langerin S-ECD surface shows its considerable contribution to the overall binding affinity (Figure 7B (b)). AT 250 nM **3.0** participates up to a third of the binding. Thus, the Langerin S-ECD surface actively interacts with the naked scaffold **3.0** which is marginally recognized by DCSIGN. The non-specific binding difference between both lectinsurfaces can illustrated clearly the importance of the scaffold choice as multivalent platform targeting a C-type lectin receptor. Moreover, the contribution of unspecific binding of the scaffold is more or less relevant according to the studied CLR and has to be considered.

As a control, direct binding of the glycomimetic **D** over a Langerin S-ECD was investigated (Figure S27). The representative sensorgram shows that reference-substracted signals R_eq_, observed for high concentrations (> 125 µM), are largely below the theoretical signal R_max_ (67 RU). Thus, no K_D_ could be extracted. This result confirms the very weak affinity of the ligand D for Langerin at monovalent scale. However, the multivalent displaying of ligand **D** in **3.D** drastically improves the affinity toward Langerin (Figure 7B) and reduces the selectivity for DC-SIGN to one order of magnitude (18 nM for DC-SIGN, 205 nM for Langerin). The selectivity of ligand D do not seem transposable from the monovalent to the multivalent scale due to the avidity generated by the spatial organization emanating from scaffold **3.0.** This finding was unexpected even if the binding analysis here is delicate to interpret due to the scaffold contribution to the binding. Cluster **3.D** illustrates that avidity can transform an imperfect ligand into a potent multivalent binder.

### Binding analysis of fucoclusters based thiacalixarene 5 and 6, and calixarene 7

In this part, another class of glycoconjugates was studied. The architectural organization of thiacalixarene displaying 4 carbohydrate-based ligands (Figure 8) is considered as potent inhibitors of DC-SIGN in the case of Ebola infection assays. (13) The IC_50_ values (Table 2), extracted from a SPR inhibition assay, showed a valuable relative affinity for DC-SIGN (IC_50_ values from 17 to 26.5 µM) with only 4 fucose-derivative ligands. However, no significant difference is observed between two different topologies of calixarenes (compound 5 (Thiacalix) and 7 (Calix)). This observation could result from the fact that their affinities are closed to the affinity limit of the reported system. In order to eliminate suspicion of limitation regarding their previous affinity estimated by competition assays (Table 2), compound 5, 6 and 7 were titrated over a DC-SIGN S-ECD surface (Figure S28) and K_D app_ are summarized in Table 2.

**Table 2:**
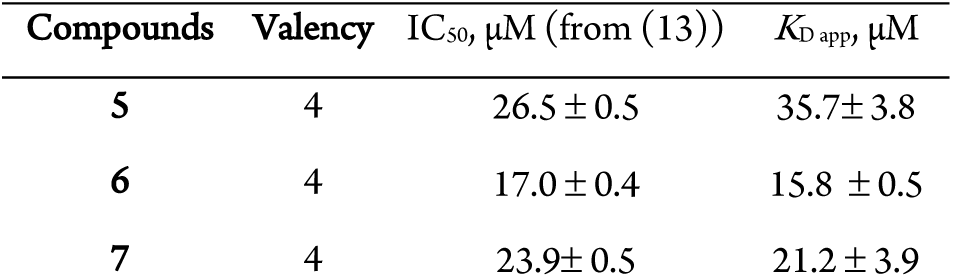
IC_50_and *K*_D app_ values of DC-SIGN inhibition for compound **5**, **6**, **7**, using respectively SPR assay and direct interaction.

**Figure 8.**
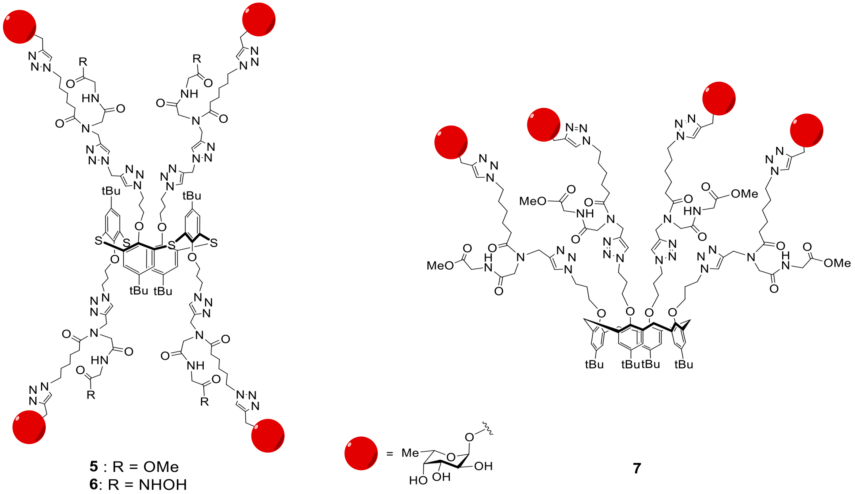
Thiacalixarene fucocluster structures.

One interesting finding is that K_D app_ values are comparable to the IC_50_ values in that case, confirming the initial affinity estimated for these compounds. The potential avidity generated by a DC-SIGN S-ECD surface does not improve the affinity of fucoclusters for DC-SIGN.

These results suggest that the relative affinities by competition assay and direct interaction are similar for compounds with a moderate valency (4 here) and an affinity higher than the reporter system (more than 5 µM). However, for compounds with an affinity below to 10 µM (case of glycoclusters) and able to display higher level of multivalency, the affinity determination should be exclusively determined by direct interaction over an oriented DCSIGN S-ECD surface. Indeed, displaying **4** fucoses, these compounds have probably access to the same level of multivalency when DC-SIGN is in solution (13) and onto the surface. On the contrary, compounds with scaffolds **3** and **4**, with **16** ligands, fully benefit from the surface avidity effect.

## CONCLUSION

The multivalent interaction in glycosciences is considered as one of the more complex interaction due to the unpredictable avidity generated by carbohydrate multi presentation. In this paper, DCSIGN S-ECD oriented surface, that can be extended to other Ctype lectins, as shown with Langerin, were designed, characterized and validated as method to evaluate reliable affinities for promising multivalent immunomodulators. This direct interaction approach, that could mimic the real cell surface situation, is a great help to qualitatively compare different classes of promising glycoconjugates targeting DC-SIGN. The tested glycocluster library highlighted the potency of glycocluster **3D** that reached remarkable affinity with 16 glycomimetics **D**.

More broadly, these finding have suggested that the multivalent carbohydrates-lectin interactions are driven mostly by avidity transforming even a poor affinity ligand into a multivalent efficient binder. This study confirms the surface avidity concept described in Munoz et al. (30) However, while they evaluated this phenomenon using ConA lectin as a model, which is a soluble globular lectin, here we used DC-SIGN, a lectin with a very strong structural asymmetry, which is physiologically a receptor embedded in the membrane. Thus, the new oriented functionalization was essential to mimic the presentation at the cell-surface. Here, these surfaces are not anymore just a case study as it was with ConA, (30) but a simulation of a real biological situation and a real tool for screenings of antagoniste in a more realistic set up. Further improvement of these DC-SIGN surfaces may come from the functionalization level based on cell surface density observed. Indeed, depending on the DC development stages, DC-SIGN is clustered in microdomain (immature DCs) or evenly distributed on the cell surface (intermediate DCs). (17, 18) These physiological difference in organization impacts strongly on the virus recognition capabilities. The perspective to simulate such situations of cell surface density of DC-SIGN is of particular interest to analyse further, at the molecular level, on a controlled system the interaction properties of DC-SIGN-dependent viruses. This will correspond to future development of these surfaces. Finally, these surfaces conserving the tetramers organisation as well as enabling DC-SIGN receptor proximity, they will be critical tools to understand the contribution of each avidity binding modes *i.e.* chelation, clustering and rebinding effects in the overall binding. Using this direct interaction as a reliable tool, investigations with glycoconjugates with limited valencies, will therefore be explored in order to elucidate the secret of avidity generation.

## EXPERIMENTAL SECTION

### Glycocluster synthesis

For the synthesis of glycoclusters **3**.**A-D** and **4.A-C** see Supplementary Information.

### DC-SIGN S-ECD expression and purification

The cDNA containing the aminoacids 66-404 of DC-SIGN was inserted into pASK-7plus plasmid with BsaI digestion. Then, cDNA fragment including corresponding extracellular domain (ECD) of DC-SIGN, Xa cleavage site and *Strep*TagII sequence in its N-terminus part were digested with XbaI and Hind III enzymes and inserted into pET-20b plasmid (Novagen). The over-expression in Rosetta (DE3) *E.coli* strain and proteins production in inclusion bodies is described as *Tabarani et al.*(26) The functional protein was purified, first, by a *Strep*Tactin column (GE Healtcare) equilibrated in 25 mM Tris-HCl pH8, 150 mM NaCl, 4 mM CaCl_2_ (buffer A) and eluted with buffer A supplemented by 2.5 mM desthiobiotin (IBA). This step was followed by an affinity chromatography on Mannan-agarose (Sigma) and Superose-12 columns as reported in *Tabarani et al.*(26)

### Langerin S-ECD expression and purification

The cDNA containing the aminoacids 65-328 of Langerin were inserted into a pASK-6 plasmid by BsaI digestion. The cDNA sequence, corresponding to Langerin ECD sequence, Xa factor site and StreptagII sequence in its N-terminus are digested by HindIII and BamHI and then inserted into a pET-20b plasmid (Novagen). The over-expression in BL21 (DE3) *E.coli* strain and proteins production in inclusion bodies was described as previously.(31) Then, Langerin S-ECD was purified by a *Strep*Tactin column (GE Healtcare) equilibrated in 25 mM Tris-HCl pH8, 150 mM NaCl, 4 mM CaCl_2_ (buffer A) and eluted with buffer A supplemented by 2.5 mM desthiobiotin (IBA). Then, the functional Langerin was purified by affinity chromatography (Mannan-agarose).

### SPR competition assays

Surface plasmon resonance (SPR) experiments were performed on a Biacore 3000 using a CM4 chip. All of procedures regarding to surface immobilization, compound titration and binding analysis are described in the Supplementary Information.

### Lectin S-ECD surface functionalization and compound titration

SPR experiments were performed on a Biacore T200 using a CM3 chip, functionalized at 5 *μ*L/min. *Strep*Tactin (IBA company) and then lectins were immobilized on flow cells using amine-coupling method. Flow cell (Fc) Fc1 and Fc3 were prepared as reference surface. Fc1 to 4 were activated with 50 *μ*L of a 0.2M EDC/ 0.05 M NHS mixture. After this step, Fc1, Fc2, Fc3 and Fc4 were functionalized with 170 µg/mL *Strep*Tactin, and then remaining activated groups of all cells were blocked with 80 *μ*L of 1 M ethanolamine. After blocking, the four Fc were treated with 5 *μ*L of 10 mM HCl to remove no-specific bound protein and 5 *μ*L of 50 mM NaOH/ 1M NaCl to expose surface to regeneration protocol. Finally, an average of 2300 RU of *Strep*Tactin was immobilized on each surface. This procedure was repeated for the functionalization of DC-SIGN S-ECD (Fc2, 2278 RU) and Langerin SECD (Fc4, 2328 RU). Different DC-SIGN S-ECD immobilization levels have been tested (252 RU, 1340 RU and 2278 RU) and data, presented here, are extracted from the highest DC-SIGN S-ECD density for its reproducibility and its surface stability.

For direct interaction studies, increasing concentrations of compound were prepared in a running buffer composed of 25 mM Tris pH 8, 150 mM NaCl, 4 mM CaCl_2_, 0.05% P20 surfactant, and 85 μL of each sample was injected onto the surfaces at 30 μL/min flow rate. The resulting sensorgrams were reference surface corrected. The apparent affinity of compounds was determined by fitting the *steady state affinity model* (eq.1) to the plots of binding responses versus concentration.

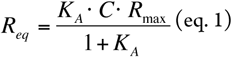

R_eq_ is equilibrium binding response, K_a_ equilibrium association constant (K_D_= 1 / K_A_), C concentration of the injected analyte and R_max_ surface binding capacity.

*K*_*D app*_ obtained reflects the affinity for the surface and not for individual lectin receptor. However, this mode of multisite interaction onto a surface is closer to the real interaction mode at the cell surface.

## ASSOCIATED CONTENT

### Supporting Information

The Supporting Information is available free of charge on the Publications website.

### Notes

The authors declare no competing financial interest.

## ACKNOWLEDGMENTS

This work used the platforms of the Grenoble Instruct center (ISBG; UMS 3518 CNRS-CEA-UGA-EMBL). SPR and MP3 platforms support from FRISBI (ANR-10-INSB-05-02) and GRAL (ANR-10-LABX-49-01) within the Grenoble Partnership for Structural Biology (PSB). This work was supported with funds from CM1102 COST Action. We are grateful to EU ITN Marie-Curie program (CARMUSYS - Grant no. 213592) for funding I. Sutkeviciute. V. Porkolab was supported by la Région Rhône-Alpes.

