## Supplementary Information for "Development of c-type lectin oriented surfaces for high avidity glycoconjugates: towards mimicking multivalent interactions on the cell surface"

ORCID ID:

1. Synthesis of glycodendrimers and characterization
2. Validation of DC-SIGN S-ECD surface
3. Binding analysis of BSAMan over a DC-SIGN S-ECD surface
4. SPR inhibition assays: sensorgrams of DC-SIGN titration over a BSAMan titration and inhibition curves of glycodendrimers
5. Binding analysis of glycodendrimers onto an oriented DC-SIGN surface. Sensorgrams and $K_{Dapp}$ determination by *Steady state affinity* model.
6. Direct interaction of glycomimetic **D** over a Langerin S-ECD surface
7. Titration of thiacalixarene fucoclusters onto a DC-SIGN S-ECD surface by SPR direct interaction.

1. Synthesis of glycodendrimers and characterization

1.1. Synthetic procedures for glycodendrimers **3.B**, **3.C**, **3.D**, **4.B** and **4.C**.

General procedures

All chemical reagents were purchased from Aldrich (Saint Quentin Fallavier, France) orAcros (Noisy-Le-Grand, France). All protected amino acids and Fmoc-Gly-Sasrin® resin was obtained from Advanced ChemTech Europe (Brussels, Belgium). For peptides and glycopeptides, analytical RP-HPLC was performed on Waters system equipped with a Waters 2695 separations module and a Waters 2487 Dual Absorbance UV/Visible Detector. Analysis was carried out at 1.23 mL/min (EC 125/3 nucleosil 300-5 C18) with UV monitoring at 214 nm using a linear A–B gradient (buffer A: 0.09% CF3CO2H in water; buffer B: 0.09% CF3CO2H in 90% acetonitrile). Purifications were carried out at 22.0 mL/min (VP 250/21 nucleosil 100-7 C18) with UV monitoring at 214 nm and 250 nm using a linear A–B gradient.

$^{1}$H and $^{13}$C NMR spectra were recorded on BrukerAvance 400 MHz spectrometer and chemical shifts (δ) were reported in parts per million (ppm). Spectra were referenced to the residual proton solvent peaks relative to the signal of CD3OD (δ 3.31 and 49.0 ppm for $^{1}$H and $^{13}$C), assignments were done by GCOSY, DEPT-135° and HSQC experiments. Standard abbreviations s, d, t, dd, bs, m, refer to singlet, doublet, triplet, doublet of doublet, broad singlet, multiplet. ESI$^{+}$-MS spectra were recorded on ThermoFischer LCQ apparatus, at university of Milano or on Waters Acquity UPLC-MS equipped with a SQ Detector 2. HRMS spectra were measured on a Waters Xevo G2-S QTof at Mass Spectrometry facility, PCN-ICMG of Grenoble. MALDI-TOF MS were performed on a Autoflex (Bruker Daltonics) using sinapinic acid matric (Sigma, 10 mg/mL in acetonitrile/water-0.1% TFA 50:50) at the mass spectrometry plateform of Institut de Biologie Structurale of Grenoble.

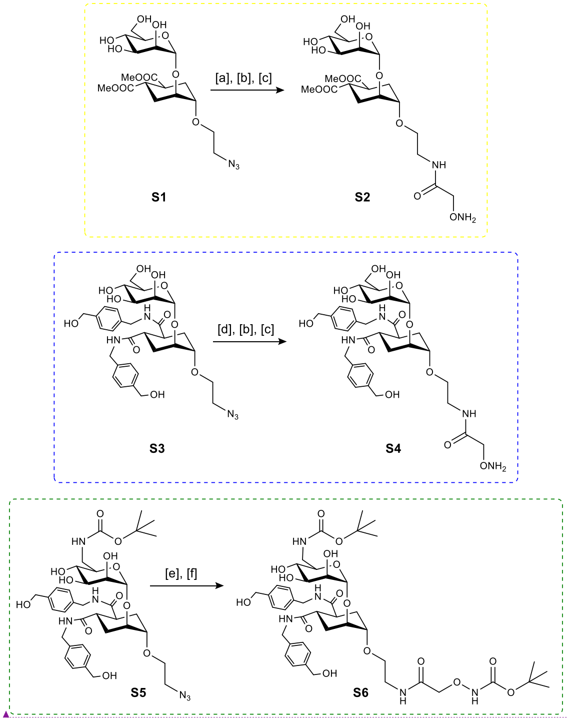


**Scheme S1.** Synthesis of mannose mimics (ligand **B**, ligand **C** and ligand **D**, see Figure 5) equipped with aminooxy moieties **S2**, **S4** and **S6**. Conditions: [a] TFA, Pd/C, $H_2$, MeOH; [b] Boc-aminooxyacetic acid *N*-hydroxysuccinimide ester, DIPEA, DMF; [c] TFA:$CH_2Cl_2$:$NH_2OH$ (60:38:2, v/v/v); [d] Lindlar-Pd, $H_2$, MeOH; [e] 1,3-propanedithiol, TEA, MeOH; [f] Boc-aminooxyacetic acid *N*-hydroxysuccinimide ester, DMF.

Code de champ modifié

Compound **S2**

To a solution of **S1**[1] (98.9 mg, 213 µmol) in freshly distilled MeOH (10 mL), under anhydrous conditions, TFA (19 µL, 248 µmol) was added. After 10 min stirring at room temperature, a catalytic amount of Pd/C was added, and the suspension was stirred under a $H_2$ atmosphere for 45 minutes. TLC ($CH_2Cl_2$:MeOH:$H_2O$ 8:2:0.25 v/v/v) revealed consumption of the starting material. The reaction mixture was then filtered through a Celite pad, washed with MeOH, concentrated under vacuum to give the corresponding amine as TFA salt. The reaction crude was used for the next reaction without any purification. ESI$^+$-MS m/z: calcd for $C_{18}H_{31}NO_{11}Na$ $[M+Na]^+$: 460.2, found: 460.5. The crude was dissolved in dry DMF (10 mL) containing DIPEA (45 µL, 287 µmol), then Boc-aminooxyacetic acid *N*-hydroxysuccinimide ester[2] (92.2 mg, 320 µmol) was added, and the reaction was stirred at room temperature for 30 minutes. The reaction mixture was concentrated under reduced pressure to dryness, then 10 mL of a TFA:$CH_2Cl_2$:$NH_2OH$ (60:38:2, v/v/v) cocktail were added. The reaction was stirred for 1 hour, then solvent mixture was concentrated and ice-cold Et2O (20 mL) was added to the residue to give a white precipitate. After centrifugation, the precipitate was washed with Et2O (10 mL), dried, then purified by preparative RP-HPLC (0-30% buffer B in 30 min.) to give pure compound **S2** (67.8 mg) as white powder after lyophilisation, in a 62% yield over three steps. $^{1}$H NMR (400 MHz, $CD_3OD$): δ 4.931 (1H, d, J = 1.3 Hz, H-1'), 4.446 (2H, bs, -COCH2ONH2), 3.988 (1H, d, J = 2.6 Hz, H-2), 3.833-3.875 (2H, m, H-2', H-6'a), 3.511-3.694 (13H, m, 2 x -CH3, -OCH2-, H-1, H-3', H-4', H-5', H-6'b), 3.443 (2H, t, J = 5.3 Hz, -CH2NH-), 2.815-2.936 (2H, m, H-4, H-5), 2.023-2.105 (2H, m, H-6b, H-3a),1.738-1.825 (2H, m, H-6a, H-3b) ppm. $^{13}$C NMR (100 MHz, $CD_3OD$): δ 176.993, 176.967 (2 x C=O), 100.419 (H-1'), 75.664, 75.638 (C-5', C-1), 73.086 (-COCH2ONH2), 72.521, 72.436, 72.399 (C-2, C-2', C-3'), 68.666 (C-4'), 68.315 (-OCH2-), 63.096 (C-6'), 52.436 (2 x -OCH3), 40.406, 40.362, 40.253 (C-4, C-5, -CH2NH-), 28.900 (C-6), 28.823 (C-3) ppm. HRMS (ESI-TOF) *m/z* calcd. for $C_{20}H_{35}N_2O_{13}$ [M+H]+: 511.2139, found: 511.2146 (error = +1.4 ppm). Analytical RP-HPLC: $t_R$ = 6.72 min (C18, λ = 214 nm 0-30% B in 15 min).

Compound **S4**

To a solution of **S3**[3] (113 mg, 168 μmol) in freshly distilled MeOH (17 mL), under anhydrous conditions, TFA (15 μL, 196 μmol) was added. After 10 min stirring at room temperature, 350 mg of Lindlar-Pd were added, and the suspension was stirred overnight under a $H_2$ atmosphere. The reaction was monitored through TLC analysis ($CHCl_3$:MeOH 7:3 v/v). The reaction mixture was then filtered through a Celite pad, washed with MeOH, concentrated under vacuum to give the corresponding amine as TFA salt. The reaction crude was used for the next reaction without any purification. $ESI^+$-MS m/z: calcd for $C_{32}H_{46}N_3O_{11}$ $[M+H]^+$: 648.3, found: 648.4; calcd for $C_{32}H_{45}N_3O_{11}Na$ $[M+Na]^+$: 670.3, found: 670.5. The crude was dissolved in dry DMF (10 mL) containing DIPEA (40 μL, 230 μmol), then Boc-aminooxyacetic acid *N*-hydroxysuccinimide ester (72.6 mg, 252 μmol) was added and the reaction was stirred at room temperature for 30 minutes. The reaction mixture was concentrated under reduced pressure to dryness, then 10 mL of a TFA:$CH_2Cl_2$:$NH_2OH$ (60:38:2, v/v/v) cocktail were added. The reaction was stirred for 1 hour, then solvent mixture was concentrated and ice-cold $Et_2O$ (20 mL) was added to the residue to give a white precipitate. After centrifugation, the precipitate was washed with $Et_2O$ (10 mL), dried, then purified by preparative RP-HPLC (0-30% buffer B in 30 min.) to give pure compound **S4** (80.5 mg) as white powder after lyophilisation, in a 66% yield over three steps. $^{1}$H NMR (400 MHz, $CD_3OD$): δ 7.204-7.287 (8H, m, Ar-H), 4.930 (1H, d, J = 1.45 Hz, H-1'), 4.560 (4H, bs, CONH-CH2-Ar), 4.414 (2H, bs, -COCH2ONH2), 4.290 (4H, bs, Ar-CH2-OH), 4.290 (4H, bs, Ar-CH2-OH), 4.003 (1H, d, J = 2.9 Hz, H-2), 3.652-3.730 (4H, m, H-5', H-3', H-6'b, 1 x -OCH2-), 3.551-3.596 (3H, m, H-1, H-4', 1 x -OCH2-), 3.448 (2H, t, J = 5.2 Hz, -CH2NH-), 2.841-2.931 (2H, m, H-4, H-5), 1.903-1.944 (4H, m, H-6a, H-6b, H-3a, H-3b) ppm. $^{13}$C NMR (100 MHz, $CD_3OD$): δ 176.987, 176.947 (2 x C=O), 141.605, 141.597, 139.020, 138.973 (4 x CAr), 128.409, 128.378, 128.176, 128.170 (8 x

CHAr), 100.237 (C-1'), 75.956 (C-3'), 75.528 (C-1), 73.246 (-COCH2ONH2), 72.572, 72.373, 72.302 (C-2, C-2', C-3'), 68.824 (C-4'), 68.041 (-OCH2-), 64.911 (Ar-CH2-OH), 63.111 (C-6'), 43.703 (CONH-CH2-Ar), 41.914, 41.788, (C-4, C-5), 40.445 (-CH2NH-), 29.498 (C-6), 28.860 (C-3) ppm. HRMS (ESI-TOF) *m/z* calcd. for $C_{34}H_{49}N_4O_{13}$ [M+H]+: 721.3296, found: 721.3301 (error = +0.7 ppm). Analytical RP-HPLC: $t_R$ = 4.84 min (C18, λ = 214 nm 0-30% B in 15 min).

Compound **S6**

To a solution of **S5**[4] (60 mg, 77.5 μmol) in dry MeOH (1.3 mL), 1,3-propanedithiol (78 μL, 775 μmol) and triethylamine (108 μL, 775 μmol) were added. The reaction was stirred at 40 °C overnight under nitrogen atmosphere. The formation of a white precipitate was observed. After completion of the reaction, the crude was filtered over a cotton pad, washed with MeOH, and the filtrate was dried under reduced pressure to obtain the corresponding amine, which was used for the next step without any purification. $R_f$ = 0.1 ($CH_2Cl_2$:MeOH:$H_2O$ = 8.5:1.5:0.1). $^1$H NMR (400 MHz, CD3OD) δ: 7.27 (d, J = 7.9 Hz, 4H, Ar-H); 7.22 (d, J = 7.9 Hz, 4H, Ar-H); 4.91 (br s, 1H, H-1'); 4.56 (bs, 4H, HO-C*H*$_2$-Ar-); 4.30 (bs, 4H, -CONH-C*H*$_2$-Ar-); 4.00-3.94 (m, 1H, H-2); 3.91-3.87 (m, 1H, H-2'); 3.75-3.43 (m, 7H, H-1, H-3', H-4', H-5', H-6'a, -OC*H*$_2$CH$_2$NH$_2$); 3.30-3.21 (m, 1H, H-6'b); 2.99-2.84 (m, 4H, H-4, H5, -OCH$_2$C*H*$_2$NH$_2$); 2.03-1.83 (m, 4H, H-3, H6); 1.91 (s, 9H, *t*Bu) ppm. The crude was dissolved in dry DMF (1.0 mL) and Boc-aminooxyacetic acid *N*-hydroxysuccinimide ester (33.5 mg, 116 μmol) was added, then the reaction stirred at room temperature for 1 h. After reaction completion the solvent was removed under reduced pressure, and the crude was purified by automated flash chromatography (silica, $CH_2Cl_2$ with gradient of MeOH from 0 to 15%) to afford 57 mg of **S6** as a solid in 80% yield over two steps. $R_f$ = 0.3 ($CH_2Cl_2$:MeOH:$H_2O$ = 8.5:1.5:0.1). $[\alpha]_D^{25}$: +8.5 (c = 0.99 in MeOH). $^1$H NMR (400 MHz, $CD_3OD$) δ: 7.28 (d, J = 8.0 Hz, 4H, H-Ar); 7.22 (d, J = 8.0 Hz, 4H, H-Ar); 4.91 (br s, 1H, H-1'); 4.56 (bs, 4H, HO-C*H*$_2$-Ar-); 4.31-4.25 (m, 6H, -CONH-C*H*$_2$-Ar-, -COC*H*$_2$ONHBoc); 4.02-3.96 (m, 1H, H-2); 3.90-3.86 (m, 1H, H-2'); 3.74-3.60 (m, 4H, -OC*H*$_2$CH$_2$NH-, H-3', H-1); 3.58-3.41 (m, 5H, H-4', H-5', H-6'a, -OCH$_2$C*H*$_2$NH-); 3.27-3.18 (m, 1H, H-6'b); 3.00-2.80 (m, 2H, H-4, H-5); 2.03-1.82 (m, 4H, H-3, H-6); 1.46 (s, 9H, *t*Bu); 1.43 (s, 9H, *t*Bu) ppm. $^{13}$C NMR (100 MHz, $CD_3OD$) δ: 177.04, 176.86 (*C*ONH); 171.66 (-*C*OCH$_2$ONHBoc), 159.80, 158.59 (-*C*=OO*t*Bu), 141.51, 139.05, 139.01, 128.39, 128.36, 128.14 (C-Ar); 100.69 (C-1'); 83.07, 80.21 ($C_{quat}$-*t*Bu); 76.48 (-CO*C*H$_2$ONHBoc); 76.10 (C-1); 74.05 (C-5'); 72.93 (C-2); 72.40 (C-2'); 72.26 (C-3'); 69.77 (C4'); 68.35 (-O*C*H$_2$CH$_2$NH-); 64.92 (HO-*C*H$_2$-Ar-); 43.66 (-CONH-*C*H$_2$-Ar-); 42.63 (C-6'); 41.87, 41.75 (C-4, C-5); 40.35 (-OCH$_2$*C*H$_2$NH-); 29.87, 29.20 (C-3, C-6); 28.86, 28.54 (*t*Bu) ppm. HRMS (ESI-TOF) *m/z* calcd. for $[C_{44}H_{65}N_5O_{16}Na]^+$: 942.43240; found: 942.43344 (error = +1.1 ppm).

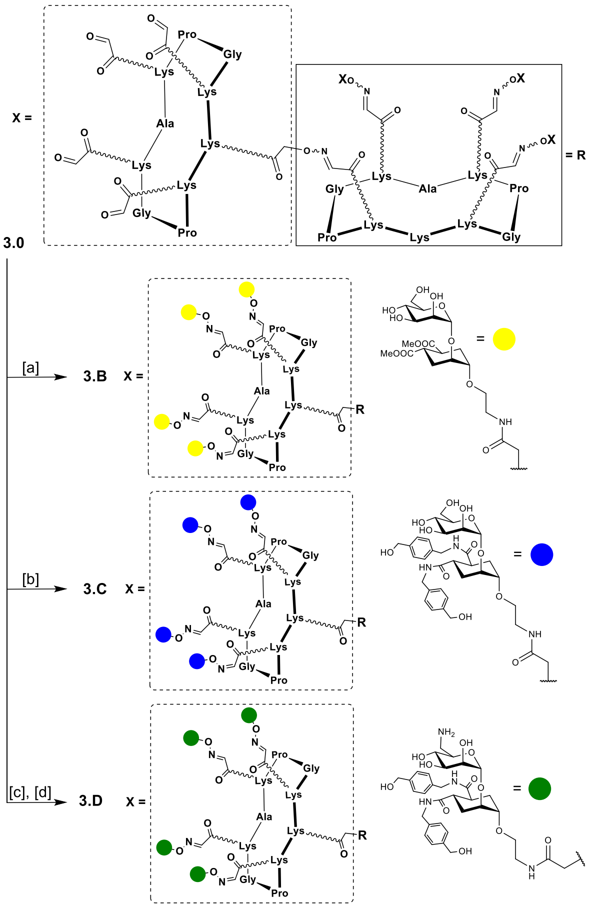


**Scheme S2.** Synthesis of mannose mimics-containing hexadecavalent glycodendrimers **3.B**, **3.C** and **3.D**. Conditions: [a] **S2**, 0.1% TFA in $H_2O$, 37 ºC; [b] **S4**, 0.1% TFA in $H_2O$, 37 ºC; [c] TFA/$CH_2Cl_2$ (1:1, v/v); [d] **S6**, 0.1% TFA in $H_2O$, 37 ºC.

Compound **3.B**

A solution of **3.0**[5] (5.2 mg, 0.81 µmol) and **S2** (12.2 mg, 23.9 µmol) in water (200 µL) containing 0.1% of TFA was heated at 37 °C for 45 minutes. The reaction mixture was directly purified through preparative RP-HPLC (5-100% buffer B in 30 min), to afford compound **3.B** (8.0 mg, 69% yield) as white powder after lyophilisation. MALDI-TOF m/z: calcd for $C_{603}H_{942}N_{111}O_{286}$ $[M+H]^+$: 14322.5 , found: 14328.6 (+6.1, error = 426 ppm). Analytical RP-HPLC: $t_R$ = 5.38 min (C18, $\lambda$ = 214 nm 5-100% B in 15 min.).

Compound **3.C**

A solution of **3.0** (3.9 mg, 0.60 µmol) and **S4** (12.1 mg, 16.8 µmol) in water (200 µL) containing 0.1% of TFA was heated at 37 °C for 45 minutes. The reaction mixture was directly purified through preparative RP-HPLC (5-100% buffer B in 30 min), to afford compound **3.C** (6.8 mg, 64% yield) as white powder after lyophilisation. MALDI-TOF m/z: calcd for $C_{827}H_{1166}N_{143}O_{286}$ $[M+H]^{+}$: 17686.9 , found: 17691.4 (+4.3, error = 254 ppm). Analytical RP-HPLC: $t_R$ = 5.13 min (C18, λ = 214 nm 5-100% B in 15 min.).

Compound **3.D**

A solution of **S6** (12.3 mg, 13.4 µmol) in a TFA/$CH_2Cl_2$ (1:1, v/v, 1.0 mL) mixture was stirred at room temperature for 30 minutes, then the solvent mixture was evaporated to dryness. The crude was added of **3.0** (4.2 mg, 0.65 µmol) and water (250 µL) containing 0.1% of TFA, then the reaction mixture at 37 °C for 45 minutes. The reaction mixture was directly purified through preparative RP-HPLC (5-100% buffer B in 30 min), to afford compound **3.D** (7.2 mg, 63% yield). $ESI^{+}$-MS m/z: calcd for $C_{827}H_{1187}N_{159}O_{270}$ $[M+6H]^{6+}$: 2946.0, found: 2947.1; calcd for $C_{827}H_{1188}N_{159}O_{270}$ $[M+7H]^{7+}$: 2525.3, found: 2526.0; calcd for $C_{827}H_{1189}N_{159}O_{270}$ $[M+8H]^{8+}$: 2209.8, found: 2210.4; calcd for $C_{827}H_{1190}N_{159}O_{270}$ $[M+9H]^{9+}$: 1964.4, found: 1964.9; calcd for $C_{827}H_{1191}N_{159}O_{270}$ $[M+10H]^{10+}$: 1768.0, found:1768.6; calcd for $C_{827}H_{1192}N_{159}O_{270}$ $[M+11H]^{11+}$: 1607.4, found: 1607.8; calcd for $C_{827}H_{1193}N_{159}O_{270}$ $[M+12H]^{12+}$: 1473.5, found: 1474.1; calcd for $C_{827}H_{1194}N_{159}O_{270}$ $[M+13H]^{13+}$: 1360.2, found: 1359.9 (mean error = 30 ppm). Analytical RP-HPLC: $t_R$ = 5.10 min (C18, λ = 214 nm 5-100% B in 15 min.).

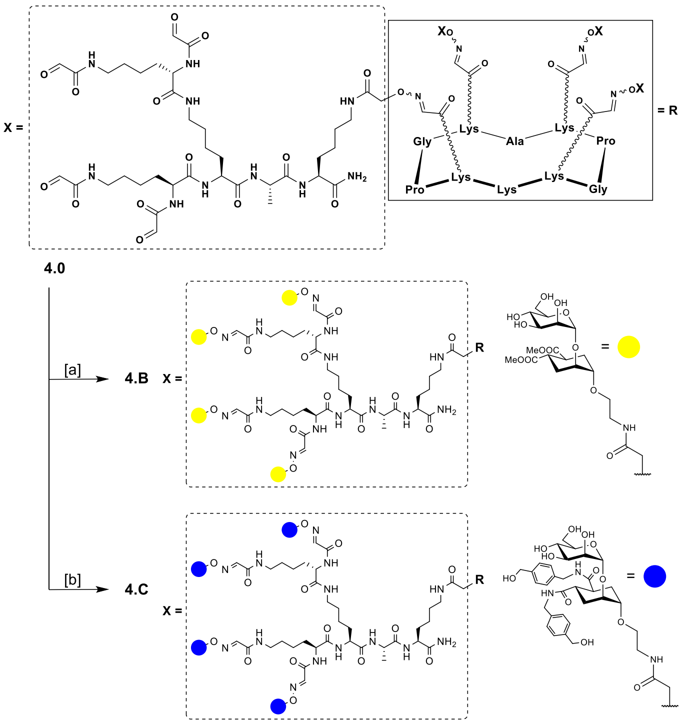


**Scheme S3.** Synthesis of mannose mimics-containing hexadecavalent glycodendrimers **4.B** and **4.C**. Conditions: [a] **S2**, 0.1% TFA in $H_2O$, 37 ºC; [b] **S4**, 0.1% TFA in $H_2O$, 37 ºC.

Compound **4.B**

A solution of **4.0**[5] (5.0 mg, 1.05 µmol) and **S2** (15.7 mg, 30.6 µmol) in water (200 µL) containing 0.1% of TFA was heated at 37°C for 45 minutes. The reaction mixture was directly purified through preparative RP-HPLC (5-100% buffer B in 30 min.), to afford compound **4.B** (8.7 mg, 66% yield) as white powder after lyophilization. MALDI-TOF m/z: calcd for C523H826N91O266 [M+H]+: 12644.6 , found: 12651.5 (+6.9, error = 546 ppm. Analytical RP-HPLC: $t_R$ = 5.26 min (C18, λ = 214 nm 5-100% B in 15 min).

Compound **4.C**

A solution of **4.0** (3.0 mg, 0.63 µmol) and **S4** (12.6 mg, 17.5 µmol) in water (200 µL) containing 0.1% of TFA was heated at 37°C for 45 minutes. The reaction mixture was directly purified through preparative RP-HPLC (5-100% buffer B in 30 min.), to afford compound **4.C** (6.9 mg, 68% yield) as white powder after lyophilisation. MALDI-TOF m/z: calcd for $C_{747}H_{1050}N_{123}O_{266}$ $[M+H]^+$: 16009.0, found: 16015.2 (+6.2, error = 387 ppm). Analytical RP-HPLC: $t_R$ = 5.01 min (C18, λ = 214 nm 5-100% B in 15 min).

Compounds **1.A**, **2.A**, **3.A** and **4.A** were synthesized according to previously reported procedures.[5]

1.2. NMR spectra.

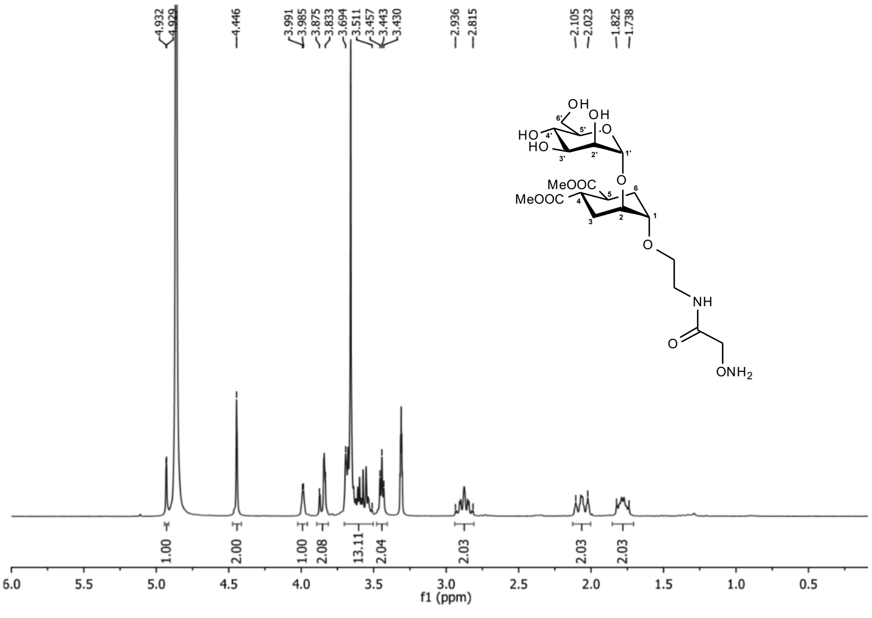


**Figure S1.** $^{1}H$ spectrum of **S2** (400 MHz, $CD_3OD$).

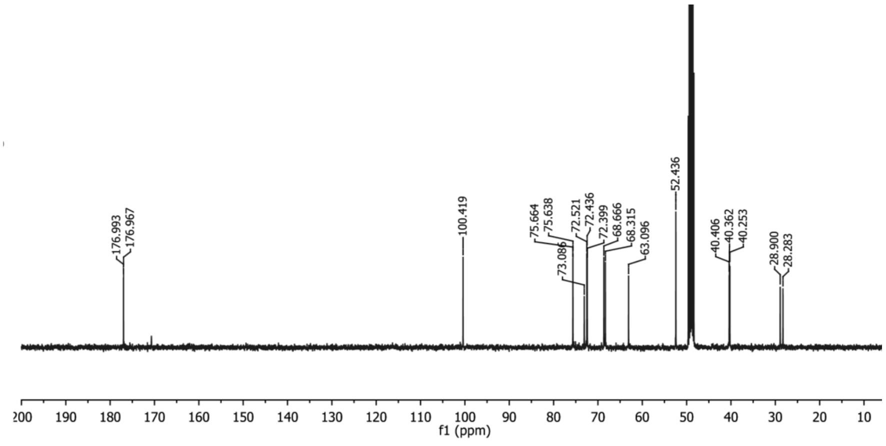


**Figure S2.** $^{13}C$ spectrum of **S2** (100 MHz, $CD_3OD$).

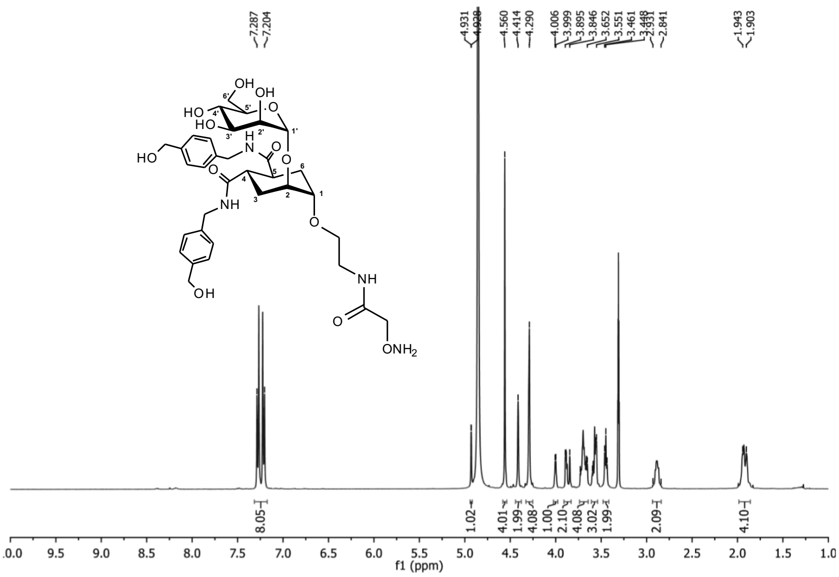

**Figure S3.** $^{1}H$ spectrum of **S4** (400 MHz, $CD_3OD$).

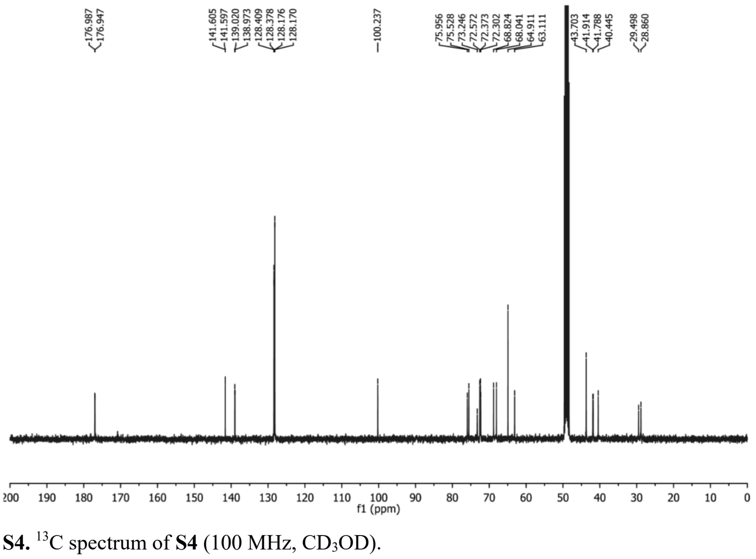

**Figure S4.** $^{13}C$ spectrum of **S4** (100 MHz, $CD_3OD$).

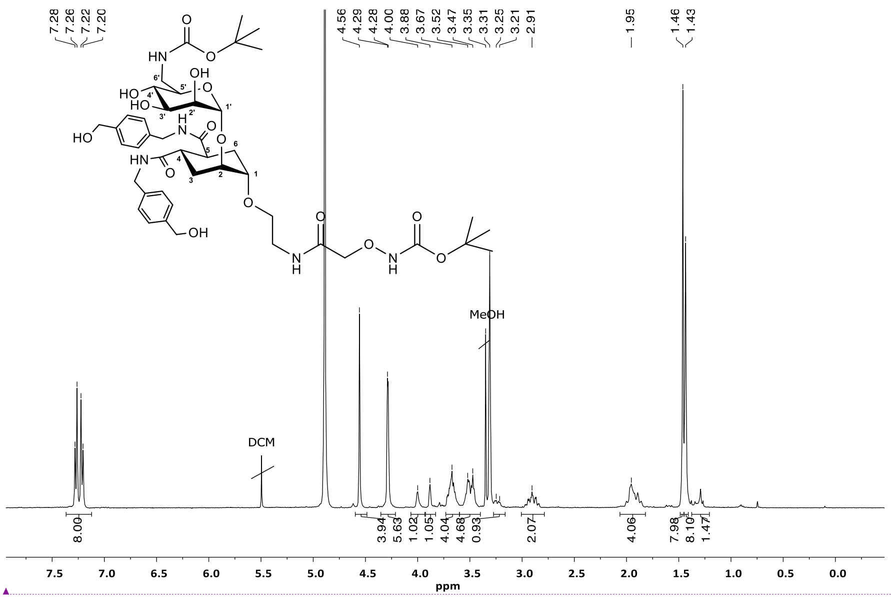


Code de champ modifié

**Figure S5.** $^{1}$H spectrum of **S6** (400 MHz, $CD_3OD$).

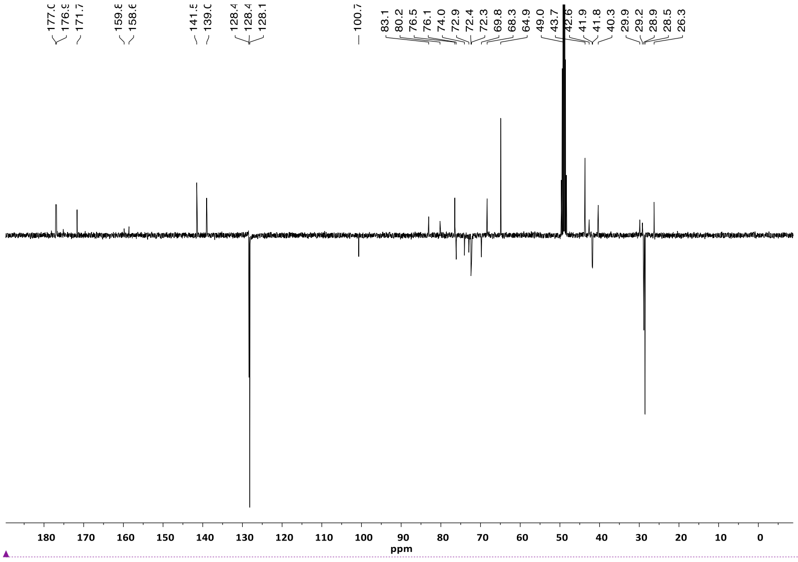


Code de champ modifié

**Figure S6.** $^{13}$C spectrum of **S6** (100 MHz, $CD_3OD$).

1.3. Mass spectra.

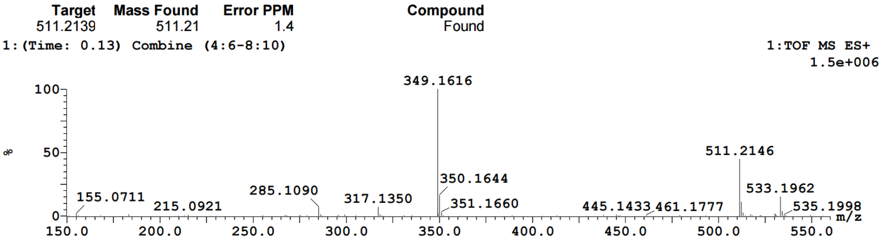


**Figure S7.** HRMS (ESI-TOF) of **S2**. *m/z* calcd for $C_{20}H_{35}N_2O_{13}$ $[M+H]^+$: 511.2139, found: 511.2146 (error = +1.4 ppm).

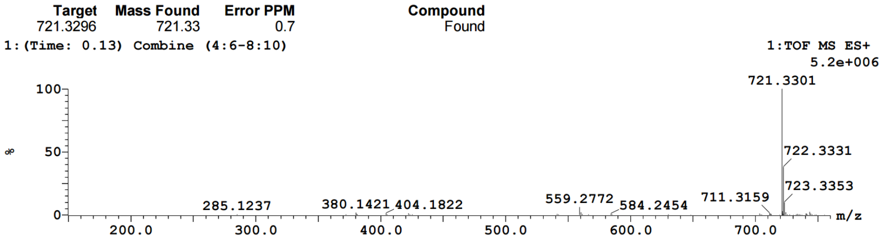


**Figure S8.** HRMS (ESI-TOF) of **S4**. *m/z* calcd for $C_{34}H_{49}N_4O_{13}$ $[M+H]^+$: 721.3296, found: 721.3301 (error = +0.7 ppm).

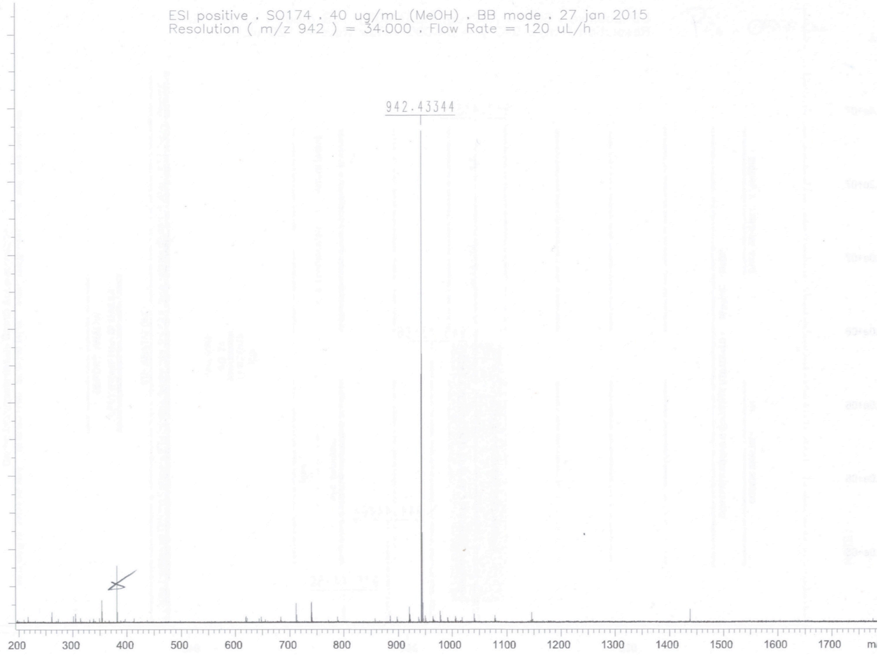


**Figure S9.** HRMS (ESI-TOF) of **S6**. *m/z* calcd. for $[C_{44}H_{65}N_5O_{16}Na]^+$: 942.43240; found: 942.43344 (error = +1.1 ppm).

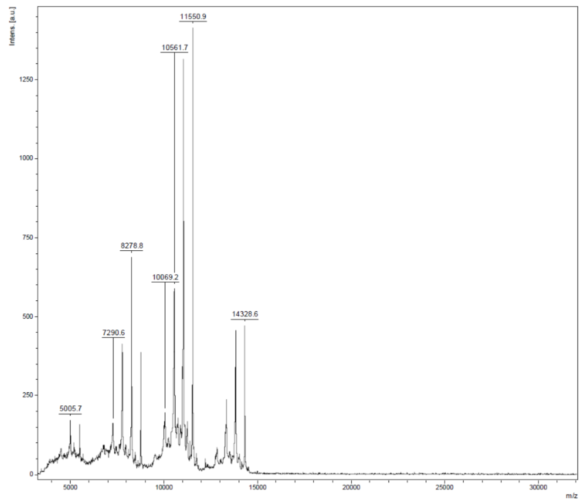


**Figure S10.** MALDI-TOF of **3.B**. *m/z* calcd for $C_{603}H_{942}N_{111}O_{286}$ [M+H]+: 14322.5 , found: 14328.6 (+6.1, error = 426 ppm).

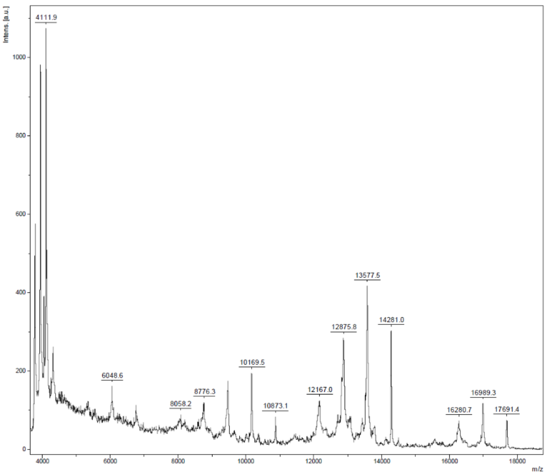


**Figure S11.** MALDI-TOF of **3.C**. *m/z* calcd for C827H1166N143O286 [M+H]+: 17686.9 , found: 17691.4 (+4.3, error = 254 ppm).

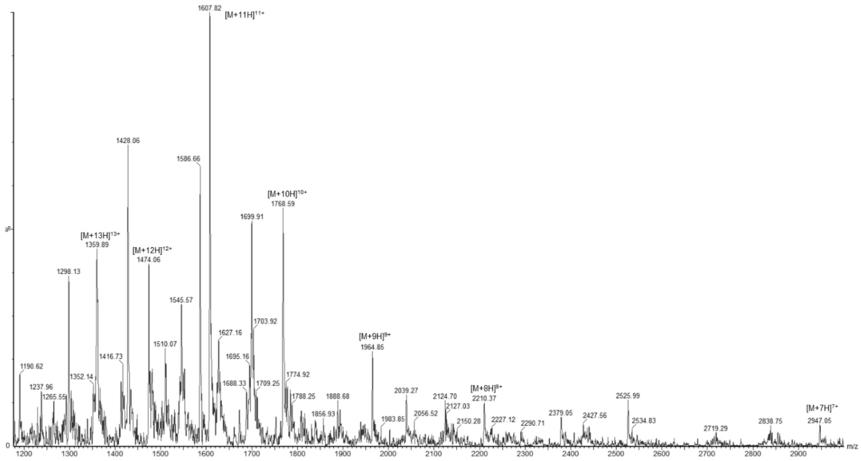


**Figure S12.** ESI$^{+}$-MS of **3.D**. *m/z* calcd for $C_{827}H_{1187}N_{159}O_{270}$ $[M+6H]^{6+}$: 2946.0, found: 2947.1; calcd for $C_{827}H_{1188}N_{159}O_{270}$ $[M+7H]^{7+}$: 2525.3, found: 2526.0; calcd for $C_{827}H_{1189}N_{159}O_{270}$ $[M+8H]^{8+}$: 2209.8, found: 2210.4; calcd for $C_{827}H_{1190}N_{159}O_{270}$ $[M+9H]^{9+}$: 1964.4, found: 1964.9; calcd for $C_{827}H_{1191}N_{159}O_{270}$ $[M+10H]^{10+}$: 1768.0, found:1768.6; calcd for $C_{827}H_{1192}N_{159}O_{270}$ $[M+11H]^{11+}$: 1607.4, found: 1607.8; calcd for $C_{827}H_{1193}N_{159}O_{270}$ $[M+12H]^{12+}$: 1473.5, found: 1474.1; calcd for $C_{827}H_{1194}N_{159}O_{270}$ $[M+13H]^{13+}$: 1360.2, found: 1359.9

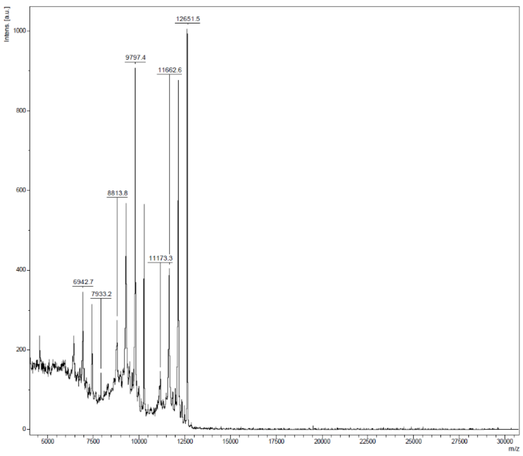


**Figure S13.** MALDI-TOF of **4.B**. *m/z* calcd for $C_{523}H_{826}N_{91}O_{266}$ $[M+H]^+$: 12644.6 , found: 12651.5 (+6.9, error = 546 ppm).

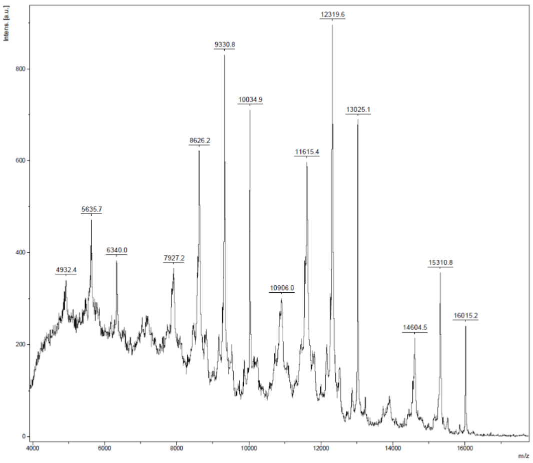


**Figure S14.** MALDI-TOF of **4.C**. *m/z* calcd for $C_{747}H_{1050}N_{123}O_{266}$ $[M+H]^+$: 16009.0, found: 16015.2 (+6.2, error = 387 ppm).

1.4. HPLC chromatograms.

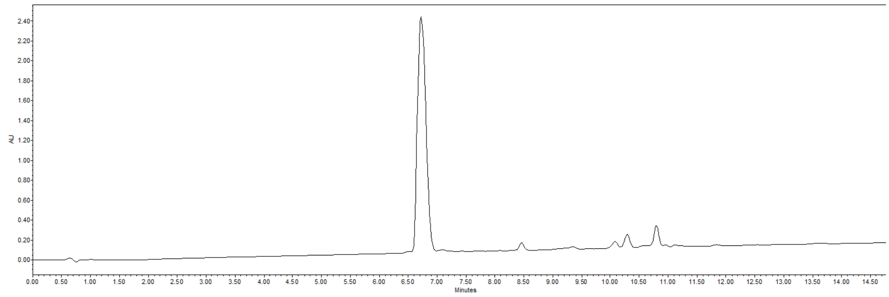


**Figure S15.** Analytical RP-HPLC of **S2**. $t_R$ = 6.72 min (C18, λ = 214 nm 0-30% B in 15 min).

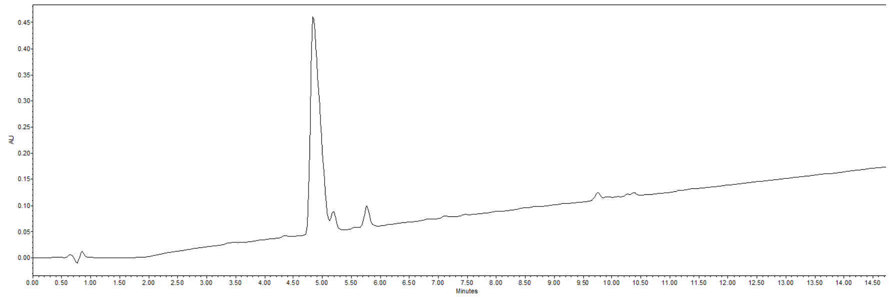


**Figure S16.** Analytical RP-HPLC of **S4**. $t_R$ = 4.84 min (C18, λ = 214 nm 0-30% B in 15 min).

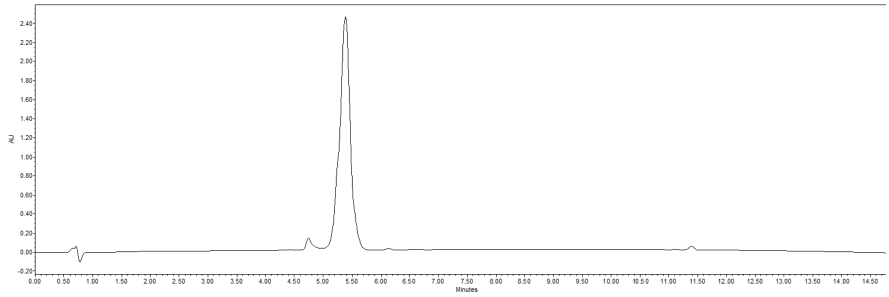


**Figure S17.** Analytical RP-HPLC of **3.B**. $t_R$ = 5.38 min (C18, λ = 214 nm 5-100% B in 15 min).

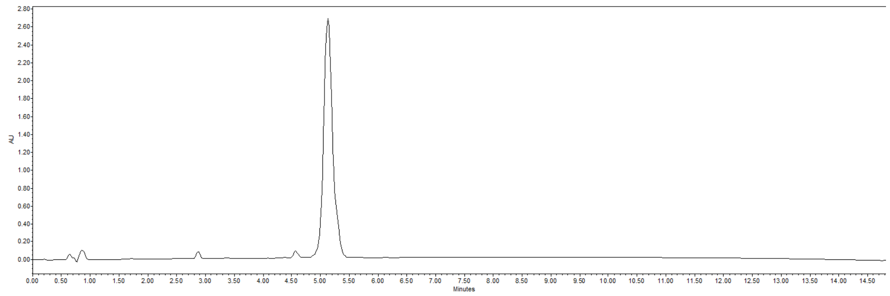


**Figure S18.** Analytical RP-HPLC of **3.C**. $t_R$ = 5.13 min (C18, λ = 214 nm 5-100% B in 15 min).

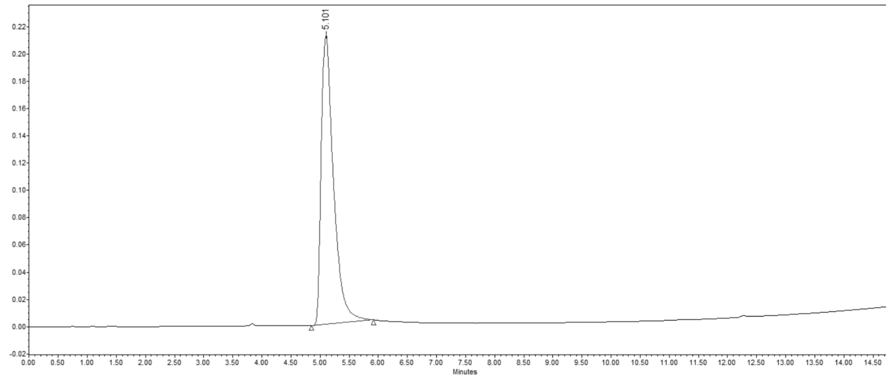


**Figure S19.** Analytical RP-HPLC of **3.D**. $t_R$ = 5.10 min (C18, λ = 214 nm 5-100% B in 15 min.).

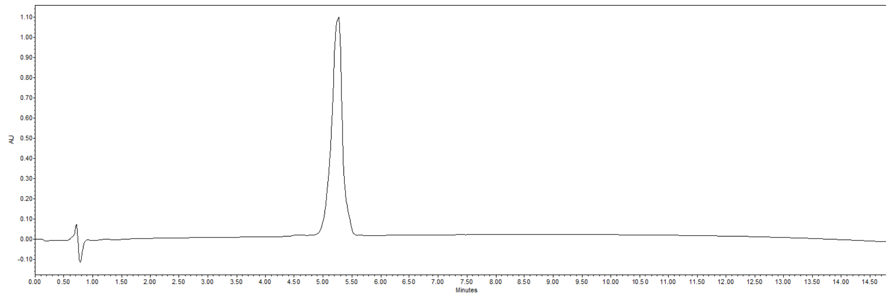


**Figure S20.** Analytical RP-HPLC of **4.B**. $t_R$ = 5.26 min (C18, λ = 214 nm 5-100% B in 15 min).

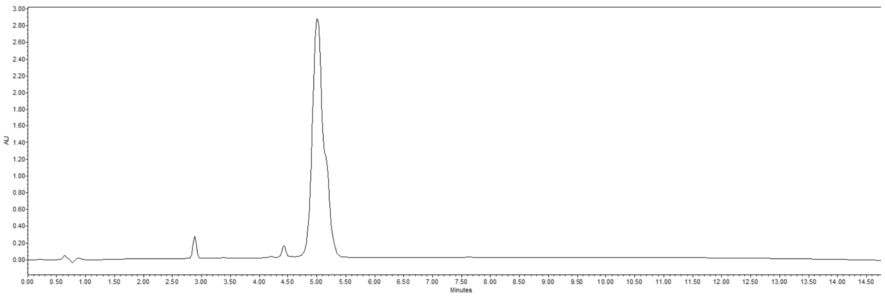

**Figure S21.** Analytical RP-HPLC of **4.C**. $t_R$ = 5.01 min (C18, $\lambda$ = 214 nm 5-100% B in 15 min).

### 1.5. Bibliography.

2. Validation of DC-SIGN S-ECD surface

2.1. DC-SIGN S-ECD affinity to StrepTactin

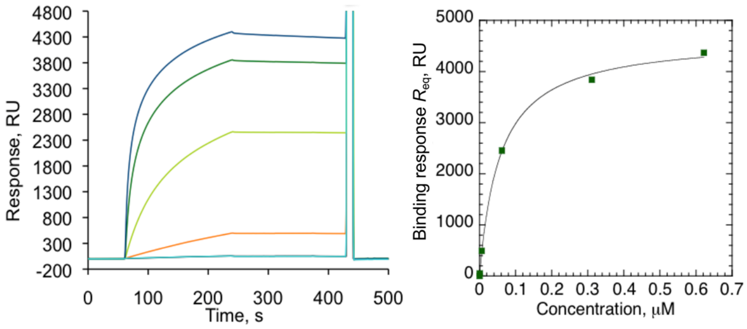


**Figure S22**. DC-SIGN S-ECD titration over a Streptactin surface. Left panel shows the sensorgram of DC-SIGN S-ECD (concentrations from 0.62 nM to 622 nM) over immobilized Streptactin (2700 RU). The right panel represents the corresponding binding responses as a function of DC-SIGN concentration. The $K_D$ extracted from *steady state affinity* model is equal to 57 nM.

2.2. Validation of DC-SIGN immobilization through StrepTag II

The dextran/*Strep*-Tactin® surface was reactivated by injection of EDC/NHS mixture. Then DC-SIGN S-ECD or DC-SIGN ECD construct prepared at 60 μg/mL concentration in HBS-P buffer were injected (150 μL/min) over reactivated surface at a flow rate of 5 μL/min of HBS-P running buffer. The remaining activated -COOH groups were blocked by 30 μL injection of 1 M ethanolamine pH 8.

2.3. Stability controls over a DC-SIGN S-ECD surface

10 consecutive injections of 0.56 μM Man-BSA over DC-SIGN S-ECD surface (3000 RU). Each injection was followed by regeneration with a mix consisting of 50 mM Gly-NaOH pH 11.9 / 0.15% TritonX100 / 25 mM EDTA pH 8, injected (8 μL) at 100 μL/min flow rate.

3. Binding analysis of BSAMan over a DC-SIGN S-ECD surface

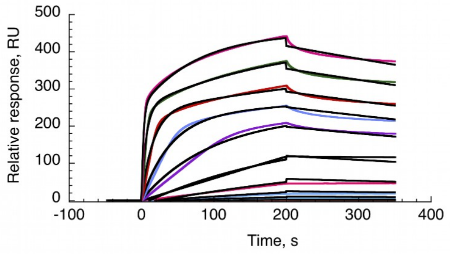


**Figure S23**: Kinetic analysis of BSAMan titration over a DC-SIGN S-ECD surface. The fit (black cruves) derives from the *heterogeneous ligand* model. The extracted values from the fit are: $K_{D1}$=6.48 nM and $K_{D2}$= 0.27 nM. The respective Rmax are 209.9 RU and 233.8 RU.

The BSAMan interaction with DC-SIGN S-ECD was evaluated first quantitatively by kinetic. As indicated, the interaction study is complex to analyse because multivalent interactions occur between 4 available CRDs of DC-SIGN and 12 glycosylated sites of BSAMan The first examined approach and the extinctive one is the *1:1 binding* fit that could be represent the avidity generation at the surface. Unfortunately, the corresponding model is not applicable in that case but also with different DC-SIGN S-ECD immobilization levels (200 RU, 1532 RU, 2300 RU). Interestingly, the *heterogeneous ligand* model is considered as relevant because the analyte could interact with 4 identical binding sites. Even if the model is limited to 2 ligand sites, this kinetic model shows a better fit than 1:1 binding (Figure S22). However, the kinetic using this model is working only with BSAMan or other glycoproteins but not with rationally designed multivalent compounds.

Regarding the *fitting steady state affinity* model, the $R_{eq}$ values (the response at e*quilibrium*) were plotted against the compound concentrations. For all of experiments, the experimental $R_{max}$ was compared to the theoretical $R_{max}$.(data not shown) and confirmed that the maximal plateau was reached even if the lowest concentrations don't reach the equilibrium at the end of association phase. We consider that the fit by a simple Langmuirian 1:1 binding model appears to be the most appropriate for regarding to the complexity of multivalent interaction.

4. SPR inhibition assays: sensorgrams of DC-SIGN - BSAMan surface binding and inhibition curves of glycodendrimers

Surface plasmon resonance (SPR) experiments were performed on a Biacore 3000 using a CM4 chip. Flow cells (Fc) 1 and 2 are functionalized at 5 µL/min. Fc were activated with 50 µL of a 0.2 M EDC/ 0.05 M NHS mixture. After this step, Fc1 and Fc2-3-4 were respectively functionalized with bovine serum albumine (BSA) and mannosylated bovine serum albumine (BSA-Man, BSA-manα1-3[manα1-6]man, Dextra laboratories, 60 µg.mL$^{-1}$). Then remaining activated groups of both cells were blocked with 30 µL of 1 M ethanolamine. After blocking, the four Fc were treated with 5 µL of 10 mM HCl to remove unspecific bound protein and 5 µL of 50 mM EDTA to expose surface to regeneration protocol. Finally, the final immobilization level of BSA and BSA-Man are respectively 1542 RU and 1478. For inhibition studies, 20 µM of DC-SIGN ECD are mixed with increasing concentrations of inhibiting compounds in a running buffer composed of 25 mM Tris pH8, 150 mM NaCl, 4 mM $CaCl_2$, 0.005% P20 surfactant. 13 µL of each sample was injected onto the surfaces at a 5 µL/min flow rate. The resulting sensorgrams were reference surface corrected.

$$y = R_{hi} - \frac{R_{hi} - R_{lo}}{1 + \left(\frac{Conc}{A_1}\right)^{A_2}} \quad (1) \qquad IC_{50} = A_1 \cdot \left( \left( \frac{R_{hi} - R_{lo}}{R_{hi} - 50} \right)^{\frac{1}{A_2}} - 1 \right) \quad (2)$$

The DC-SIGN binding responses were extracted from sensorgrams, converted to percent residual activity values (y), which were plotted against corresponding compound concentration. The 4-parameter logistic model (equation 1), available in the BiaEval software, was fitted to the plots, and the $IC_{50}$ values were calculated, from equation 2, using the values of fitted parameters ($R_{hi}$, $R_{lo}$, $A_1$ and $A_2$).

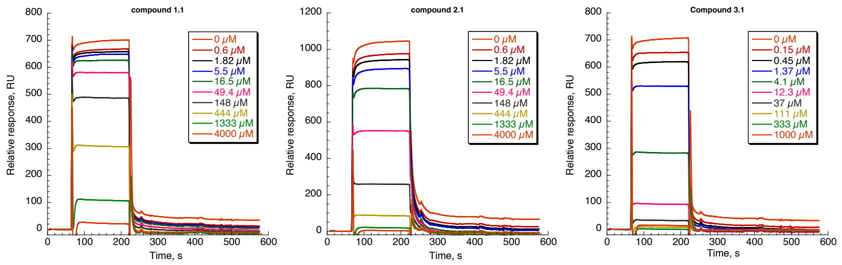

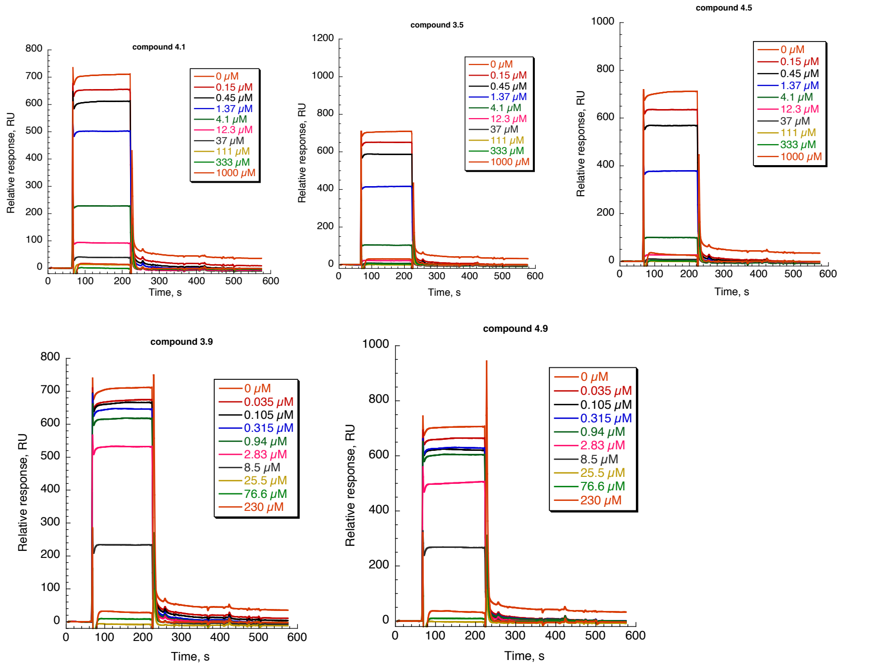


**Figure S24**: Inhibition of DC-SIGN interaction over a BSA-Man surface. 20 μM of DC-SIGN ECD and increasing concentrations of compounds were co-injected over the surface.

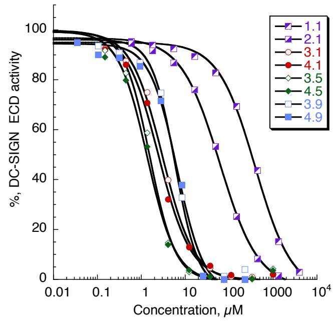


**Figure S25:** Inhibition curves of DC-SIGN interaction with BSA-Man surface. $IC_{50}$ values are extracted from the *4-parameter equation* model.

5. Binding analysis of glycodendrimers onto an oriented DC-SIGN surface. Sensorgrams and $K_{Dapp}$ determination by *Steady state affinity* model.

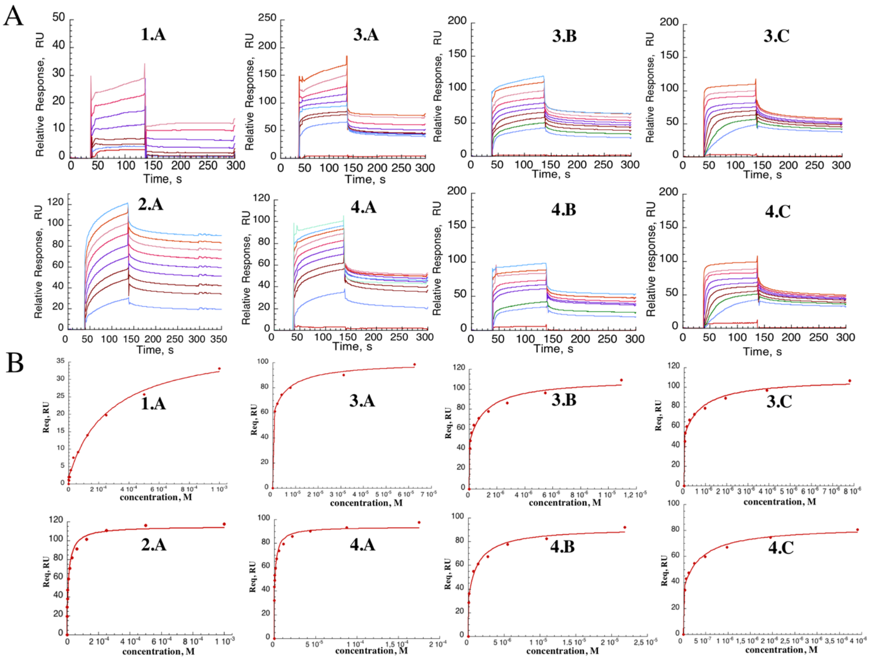


**Figure S26:** Binding analysis of glycodendrimers over an oriented DC-SIGN surface. (A) Reference surface corrected sensorgrams showing the binding of indicated compound over a DC-SIGN S-ECD surface. (B) Binding responses were plotted against their concentrations and fitted by a *steady state affinity* model.

6. Direct interaction of glycomimetic **D** over a Langerin S-ECD surface.

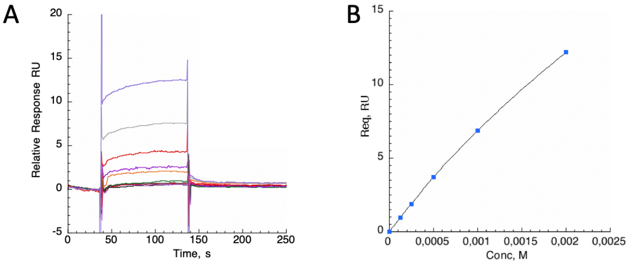


**Figure S27**: (A) Reference-subtracted sensorgram of increasing concentrations (0, 4, 8, 16, 31, 62, 125, 250, 500, 1000, 2000 µM) of glycomimetic **D** over Langerin S-ECD surface. B) Steady state binding analysis (n=1) for ligand **D** (blue dots).

7. Titration of thiacalixarene fucoclusters onto a DC-SIGN S-ECD surface by SPR direct interaction.

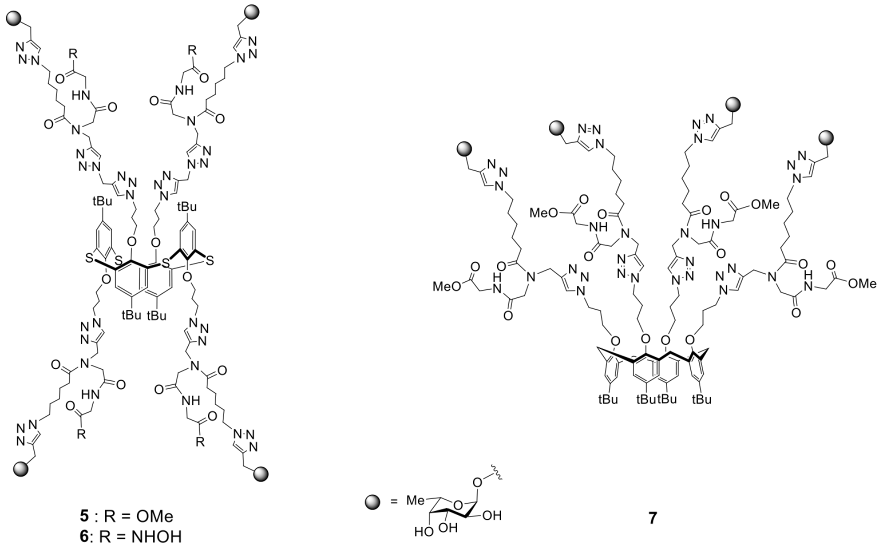


**Scheme S4 :** Thiacalixarene fucoclusters structures

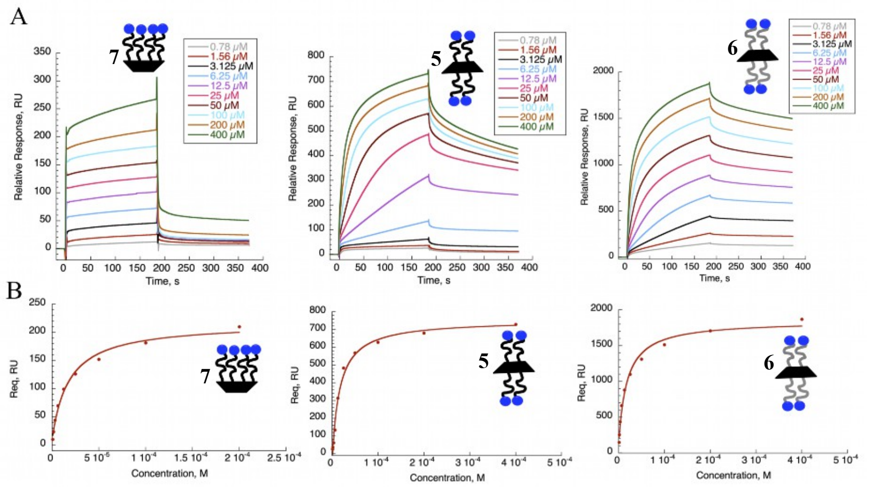


**Figure S28:** Binding analysis of thiacalixarene fucoclusters over a DC-SIGN S-ECD surface. (A) Reference surface corrected sensorgrams showing (from left to right) the binding of compound **7, 5** and **6** over a DC-SIGN surface. (B) Compound binding responses (red dots) were plotted against their concentrations. Curves (red lines) are fitted by a *steady state affinity* model (binding n=1).
